# Priority effects, consumer pressure, and soil resources independently alter plant diversity and resource strategies during a multi-year successional field experiment

**DOI:** 10.1101/722264

**Authors:** Peter A. Wilfahrt, Fletcher W. Halliday, Robert W. Heckman

**Author notes:** Corresponding author: Peter Wilfahrt.

## Abstract

- Plant community succession is structured by priority effects, plant consumer pressure, and soil resource supply. Importantly, these drivers may interact, their effects may vary temporally, and they may influence different facets of plant community diversity by promoting different plant tradeoff strategies.
- In an herbaceous successional system, we manipulated priority effects by altering initial plant richness, consumer pressure via pesticide spraying, and soil resource supply via fertilization. We examined how these processes jointly influenced succession, including taxonomic diversity and functional traits, over four years.
- Diversity decreased in different years in response to more diverse priority effects, lower consumer pressure, and increased soil resource supply. Functionally, higher soil resource supply increased community height, SLA, and seed mass; higher consumer pressure decreased intraspecific community height, and increased interspecific SLA; priority effects led to decreased seed mass only when plots were unplanted.
- Our results suggest species’ resource strategies underlie plant diversity responses. Resource addition promoted resource-acquisitive species, consumer pressure disadvantaged resource-conservative species, and diversity of priority effects altered subsequent community composition through persistence of early residents, not via traits. We show that community responses to drivers of succession depend on underlying trait tradeoffs of resident species, and these tradeoffs influence community diversity across succession.

## Introduction

The speed and direction of succession depends on co-occurring factors, including the order of species arrival (Körner *et al.*, 2008), the supply of soil nutrients (Isbell *et al.*, 2013), and the loss of tissue to plant consumers (Mordecai, 2011; Kempel *et al.*, 2015). Ecological tradeoffs often determine how individual species respond to these factors, in turn determining how these factors affect the successional trajectory of a community. These potential tradeoffs can be understood using taxonomic and functional diversity metrics, which in tandem enable inference of underlying assembly mechanisms regulating community membership (Purschke *et al.*, 2013). Ecological tradeoffs may, for instance, result in a species being a poor soil nutrient competitor but good light competitor (Dickson *et al.*, 2014), having increased susceptibility to herbivory or disease but being capable of rapid growth (Züst & Agrawal, 2017), or investing in a bet-hedging strategy enabling high dispersal of offspring to early successional and low competition patches, at the expense of high individual seedling mortality (Leishman *et al.*, 2000). The factors driving these tradeoffs, and consequently, how succession proceeds, may interact; thus considering them jointly provides additional mechanistic insight into the drivers of succession (Borer *et al.*, 2014b; La Pierre *et al.*, 2015; Heckman *et al.*, 2017). Here, by jointly studying changes in taxonomic and functional diversity over time, we identify the interactive effects of abiotic and biotic drivers and how these change temporally in an herbaceous successional system.

### Ecological factors that determine community diversity during succession

The diversity of a community early in secondary succession influences subsequent changes in diversity as early arriving species may gain an advantage through priority effects (Veen *et al.*, 2018). The succession of a recently disturbed community is characterized initially by relatively high resource availability that is reduced as species colonize and establish. A species may fail to establish in these initial periods despite being biologically suited to the environment because it either fails to disperse to that area, or it disperses and germinates but fails to persist due to competition from more mature resident species that established earlier or survived the disturbance (Tilman, 2004). The degree to which competition from residents prevents subsequently arriving species from establishing may depend on the diversity of initial residents (Fargione & Tilman, 2005). More diverse communities can more efficiently capture resources, reducing the resource availability for a new species to establish (Levine & D’Antonio, 1999).

By altering competition among species, soil resource supply and consumer presence can independently and interactively moderate the trajectory of succession. Competition for multiple limiting soil nutrients promotes species’ coexistence (Harpole & Tilman, 2007), whereas, depending on soil characteristics and traits of the species present (Suding *et al.*, 2005; Dickson *et al.*, 2014; Harpole *et al.*, 2016), increased soil resource supply leads to loss of species richness (Grime, 1973; Rajaniemi *et al.*, 2003; Harpole & Tilman, 2007; Dickson & Foster, 2011). Similarly, the presence or absence of plant consumers, such as pathogens (Mordecai, 2011) or invertebrate herbivores (Kempel *et al.*, 2015) can promote coexistence, particularly when consumers prefer dominant plants, thus promoting community evenness (Mortensen *et al.*, 2018). Conversely, plant consumer reduction may increase plant diversity when consumers more negatively impact rare species (Koerner *et al.*, 2018). These two factors may interact to alter community diversity through two pathways. First, either one or several species may be co limited by low soil resource supply and consumer presence (Lind *et al.*, 2013), allowing increased dominance when both of these factors are alleviated. Second, one or a few species may gain dominance following increased soil resource supply only in the presence of consumers, while reduced consumer pressure leads to dominance of one or a few different species (La Pierre *et al.*, 2015; Koerner *et al.*, 2018). This could prevent increases in dominance of any one species when both limiting factors are alleviated. Finally, increased soil resource supply and reduced consumer pressure may both govern successional changes by reducing light availability below the plant canopy (Borer *et al.*, 2014b), thereby restricting more quickly the light demanding species that tend to dominate early in succession.

Priority effects arising from more diverse communities may mitigate diversity losses following increased soil resource supply (Hodapp *et al.*, 2018). This may occur because increased plant richness can increase soil resource use efficiency (Hooper & Vitousek, 1998), preventing or slowing competitive exclusion by a single dominant species. However, this pattern has been observed to reverse over time (Weisser *et al.*, 2017), meaning that a persistent increase in soil resource supply could overwhelm any initial diversity effects. Consumer pressure effects may be conditioned on initial plant diversity as early community dominance of a species may be maintained by reducing consumer pressure (Olff & Ritchie, 1998). Moreover, succession is a temporal process and may result in soil resource supply or consumer pressure effects manifesting only after several years (Isbell *et al.*, 2013; La Pierre *et al.*, 2015; Weisser *et al.*, 2017). Thus, diversity underlying priority effects could moderate the rate of change in diversity during succession following changes to soil resource supply and consumer pressure. Low initial diversity, increased soil resource supply, and reduced consumer pressure could all lead rapidly to a high dominance of one or a few species, provided they favor the same species. However, diversity trajectories are less clear when these drivers impede one another or favor different species and trait-based approaches are an effective way to detect this (Purschke *et al.*, 2013).

### Mechanisms underlying diversity changes

Ecological tradeoffs across species underlie diversity changes during succession, and functional traits may clarify the prevalent mechanisms controlling these tradeoffs. Specifically, individual species’ investments in seed, height, and leaf traits (e.g., Westoby 1998) may reflect the ecological strategies of constituent species at a given time in response to shifting resource environments during succession (Webb *et al.*, 2010). Seed mass captures a competition-colonization tradeoff among species (Turnbull *et al.*, 1999; Mouquet *et al.*, 2004), where small-seeded species are adapted to a colonization strategy allowing high dispersal to open habitats, but decreased chances of survival when germinating underneath extant vegetation (Leishman *et al.*, 2000). As priority effects, plant consumers, and soil resource supply affect the availability of resources, seed mass acts as a filter for establishment. Because competition for light is asymmetric (DeMalach *et al.*, 2017), vegetative height is a straightforward trait for understanding light competition (Westoby *et al.*, 2002). To grow taller than its neighbors, an individual must invest in structural biomass, potentially at the cost of root or leaf biomass (Shipley & Meziane, 2002). Finally, specific leaf area (SLA) represents a resource conservation-acquisition tradeoff (Poorter *et al.*, 2009). An increasing amount of leaf area per energy invested (i.e. increased SLA) may increase the growth rate of a species at the cost of leaf life span (Wright *et al.*, 2004), which may be especially disadvantageous when plant consumer pressure is high (Coley *et al.*, 1985). Thus, as environmental resource limitations shift, so too do competitive hierarchies, which may be reflected in community SLA values.

Ultimately, priority effects, consumer pressure, and soil resource supply may simultaneously influence the taxonomic and functional diversity of plant communities throughout succession. Examining temporal trait responses within the same system can reveal the relative importance of these processes through time. In this study, we examine successional responses of community taxonomic diversity and LHS traits across four years in a multifactor field experiment manipulating initial plant diversity, soil resource supply, and consumer pressure. After constructing initial communities with various compositions, we allowed natural colonization and monitored community diversity for four years. We use this system to examine 1) how altered biotic and abiotic conditions influence species diversity, 2) how leaf-height-seed traits help explain diversity changes, 3) how these influences change through time as the community develops, and 4) how the abiotic and biotic conditions interact with each other during succession.

## Methods

### Study area

This study was conducted at Widener Farm, an old field maintained as part of Duke Forest Teaching and Research Laboratory in the Piedmont of North Carolina, USA. Widener Farm was used for row crops from the mid-1950s until 1996, and has since been maintained as an herbaceous community by annual mowing. The site receives an average of 1221 mm of annual precipitation. It is co-dominated by perennial grasses, *Andropogon virginicus* and *Lolium arundinaceum*, but maintains a rich diversity of annual and perennial herbaceous species (Heckman *et al.*, 2016) experiencing a range of consumer damage (Halliday *et al.*, 2017, 2019).

### Experimental design

In order to test the individual and interactive effects of priority effects, consumer pressure, and soil resource supply on community dynamics, we used a randomized, complete block design with factorially crossed treatments of each factor. In 2011, we denuded 1 × 1 m plots of vegetation by applying glyphosate herbicide (Riverdale^®^ Razor^®^ Pro, Nufarm Americas Inc, Burr Ridge, IL); two weeks later, we removed dead vegetation and covered plots with landscape fabric to impede natural recolonization. One-meter wide alleys between plots were left vegetated.

Priority effects were manipulated by assigning plots to one of three treatment levels: monoculture, five-species polycultures, and unplanted. The planted species pool were six perennial, native, herbaceous species already occurring at Widener Farm. Individuals were allowed to establish for 2011, and in 2012 we replaced all dead individuals. In July 2012, we weeded plots of all non-planted species and removed the landscape fabric without damaging planted individuals. Following this, no further weeding was done as natural colonization proceeded. Unplanted plots were prepared similarly to the other plots, but no species were planted in the denuded plots in order to quantify diversity changes during succession with no initial aboveground competition. There were six possible polyculture species combinations, each excluding one of the six planted species, six possible monocultures, and the unplanted treatment, creating 13 possible initial community compositions. These 13 compositions were factorially crossed with the soil resource supply and consumer pressure treatments and replicated once in each of five spatial blocks (n=260 plots).

Consumer pressure was manipulated by assigning plots to one of two treatment levels: control and pesticide application. We applied a fungicide and an insecticide every two to three weeks during the growing season from July 2012 to September 2015; neither pesticide showed non-target effects on plant growth of common Widener Farm species under greenhouse conditions (Heckman *et al.*, 2016). Soil nutrient supply was manipulated by assigning plots to one of two treatment levels: control and fertilization. Fertilized plots received an annual application of 10 g m^-2^ slow-release NPK (e.g, following Borer *et al.*, 2014a). The first application occurred after we removed the landscape fabric in July 2012 and in May of each subsequent year. Further details on planting and spraying can be found in the supplementary materials.

### Plant community composition

We measured plant community composition each year at peak biomass in early September by visually estimating absolute percent cover of all vascular plants (including planted and unplanted species) within a centrally located 0.75 × 0.75 m subplot in each plot to avoid edge effects. The first survey was conducted in September 2012 two months after natural colonization began following removal of the landscape fabric. An additional survey conducted in June of 2014 was used to inform trait data collection (described below) only. Light attenuation was measured immediately following plant cover surveys, using a light-ceptometer (AccuPAR LP-80, Decagon Devices Inc., USA), as the ratio of photosynthetically active radiation above the plant canopy to the ground level at two 0.8 m strips to the left and right of center in each plot (Borer *et al.*, 2014a).

### Taxonomic diversity

We calculated two measurements of diversity: species richness, and species evenness. To quantify evenness, we calculated Hill’s diversity for each plot in each year using total percent cover of each species as an abundance metric and the package *vegan* (Oksanen *et al.*, 2013) in R. Then, we divided this result by plot species richness to obtain a measure of evenness (*sensu* Tuomisto 2012). We calculated these indices for two sets of nested species data: all species present in a plot at the sampling period, and all colonizing species in a plot that were not planted as part of the initial richness treatment (thus one of our six planted species could still be counted as a colonizer in plots where it was not planted). This was done in order to assess the priority effects our planted species had on the diversity trajectories as a whole (total species count) and on the subsequently arriving species (colonizer species count).

### Trait data

Specific leaf area (SLA) was measured in July, 2014, immediately following a cover survey conducted in late June. In each plot, we selected species in descending order of percent cover until 80% of the relative vegetative cover of that plot was accounted for (Pérez-Harguindeguy *et al.*, 2013). Then, we selected ten leaves in each plot by cycling through its species list in descending order of cover. For instance, if six species accounted for >80% of the relative cover of a given plot, two leaves would be selected for each of the four most abundant species, and one leaf for each of the remaining two species. Leaves were chosen randomly from within the plot, but an effort was made not to sample from the same ramet when a species was sampled multiple times. In total, 2590 leaves were sampled across the experiment; an average of 4.5 species were selected per plot and 35 species were sampled across all plots.

We measured height immediately following the September 2014 cover survey using that data to determine species’ relative cover per plot. We measured the naturally-standing vegetative height of the tallest individual of each species included in the top 80% of relative cover in each plot. Because the variable of interest was a species’ height potential in any given plot, replication of a species occurred only across plots. This resulted in 1124 individuals being measured in 37 species across all plots, for an average of 4.3 species per plot.

Due to high variability of species’ phenology in the system and the absence of reproductive structures in many species, seed mass data were acquired from Royal Botanic Gardens Kew (2016) for the most abundant species (44 species total) in the experiment. Where multiple weights were reported, we took the mean value from all sources reported; two of these 44 species, *S. integrifolia* and *S. pinetorum*, were not present in the database, so we selected the value of their nearest phylogenetic neighbor. Because these data were not collected locally, we were unable to estimate within-species variation. However, variation in seed mass may be lower than the other traits in this study; several studies suggest that within species means of seed mass are conserved across environments (Violle *et al.*, 2009; Kazakou *et al.*, 2014). Seed mass values were log transformed at the species level to normalize the data as they ranged across four orders of magnitude.

Community weighted means (CWM) were calculated using species means for each trait as:

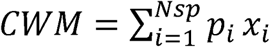

where Nsp is the number of species within a plot with a mean trait value in the dataset, p_i_ is the relative abundance of species, *i*, in the plot, and x_i_ is the species trait values. For seed mass, we calculated the CWM for each plot in each year using the year-specific relative cover value for each species that colonized a specific plot. Because our planting treatment effectively overcame dispersal limitations, we omitted any species planted in a specific plot from the relative cover and subsequent CWM calculations. As SLA and height were only measured in 2014, we only calculated CWMs for this year using the species’ relative cover as an abundance metric. For these traits, we calculated inter and intraspecific values and counted all species, whether planted or not. For SLA, we used the experiment-wide mean of each species for the interspecific value. For interspecific maximum height, we used the 95^th^ quantile value of a species’ experiment-wide measurements. This approach captures the upper end of a species’ distribution that is more representative of its height potential within the study system. To calculate intraspecific effects, we calculated a z-score for each species with 5 or more trait observations across the experiment, by centering the mean at zero and transforming the data to have a standard deviation of 1. Then, we recalculated CWMs using the observed z-score for each species in each plot; plots with multiple observations of SLA per species used the average within-plot z-score for that species. Effectively, this shows whether treatments influenced the change in traits of species relative to their own, species-intrinsic range of variation.

### Statistical analyses

All data were analyzed using R version 3.3.3 (R Core Team 2016). We used linear mixed effects models to test the effects of our treatments on four response variables across time: species richness, species evenness, colonizers’ CWM of seed mass, and light attenuation at ground level. The diversity metrics calculated with all species and colonizers only were analyzed separately. In all models, year was modeled as a categorical variable to avoid making assumptions about linearity in the temporal trajectory of community responses. Full interaction models between the three treatments and time were first tested, and non-significant interactions were sequentially dropped until the most parsimonious model was found (Zuur *et al.*, 2009). However, all treatment × year interactions were retained regardless of significance, as we were expressly interested in how treatment effects changed through time, and omission of these terms led to small, but qualitatively important changes to interpretation. Similarly, fertilization masked the effects of the other treatments in some models, so we report the results of models with all two way interactions regardless of significance in the supplementary material. We used the lsmeans package to examine the multiple comparisons of our reduced models while adjusting our p values using Tukey’s HSD (Lenth & Hervé, 2015). To account for non-independence of repeated samples in our models, we included plot as a random effect with an auto-regressive, order 1 autocorrelation structure (Zuur *et al.*, 2009). In order to account for observed heteroscedasticity between levels of the diversity treatment, we also incorporated an identity variance structure of diversity term in each model (Zuur *et al.*, 2009). We also included planted community composition as a random effect; this conservative approach allows us to ascribe observed differences between initial richness treatments to richness *per se*, rather than composition (Schmid *et al.*, 2002; Heckman *et al.*, 2017). An identity variance structure was applied to ‘year’ for the light attenuation data due to high heteroscedasticity between years. We also examined correlations between diversity metrics and light attenuation following the same procedure, comparing slope coefficients between years using the ‘lstrends’ function in the lsmeans package. Light attenuation was log-transformed to meet model assumptions. To account for spatial heterogeneity within the study, experimental blocks were included in all models as fixed effects.

## Results

### Priority effects, consumer pressure, and soil resource supply on species coexistence and dominance

We manipulated priority effects by experimentally altering the initial richness of plant communities. We hypothesized that a more diverse initial assemblage would more effectively draw down resources, limiting the number of species able to colonize a community during succession. Consistent with a priority effect, initial richness affected subsequent total richness through time (p<0.01; Fig. **1a**, Table S1, S2), as polycultures had higher total richness in the first year of the study, but no effects were observed thereafter. Furthermore, consistent with the hypothesis that more diverse initial assemblages would limit the number of species able to colonize a community, initial richness influenced the richness of colonizing species (e.g., species that were not planted, but later colonized a plot; p<0.01; Figure **1b**, Table S3) and this effect changed through time (p<0.01; Table S4). Polycultures had lower colonizing richness than monocultures or unplanted plots throughout the study, and monocultures had lower colonizing richness than unplanted plots in the first two years of study. Together, this suggests that planted species benefited from priority effects allowing them to maintain populations and limit colonizer richness.

**Fig 1.**
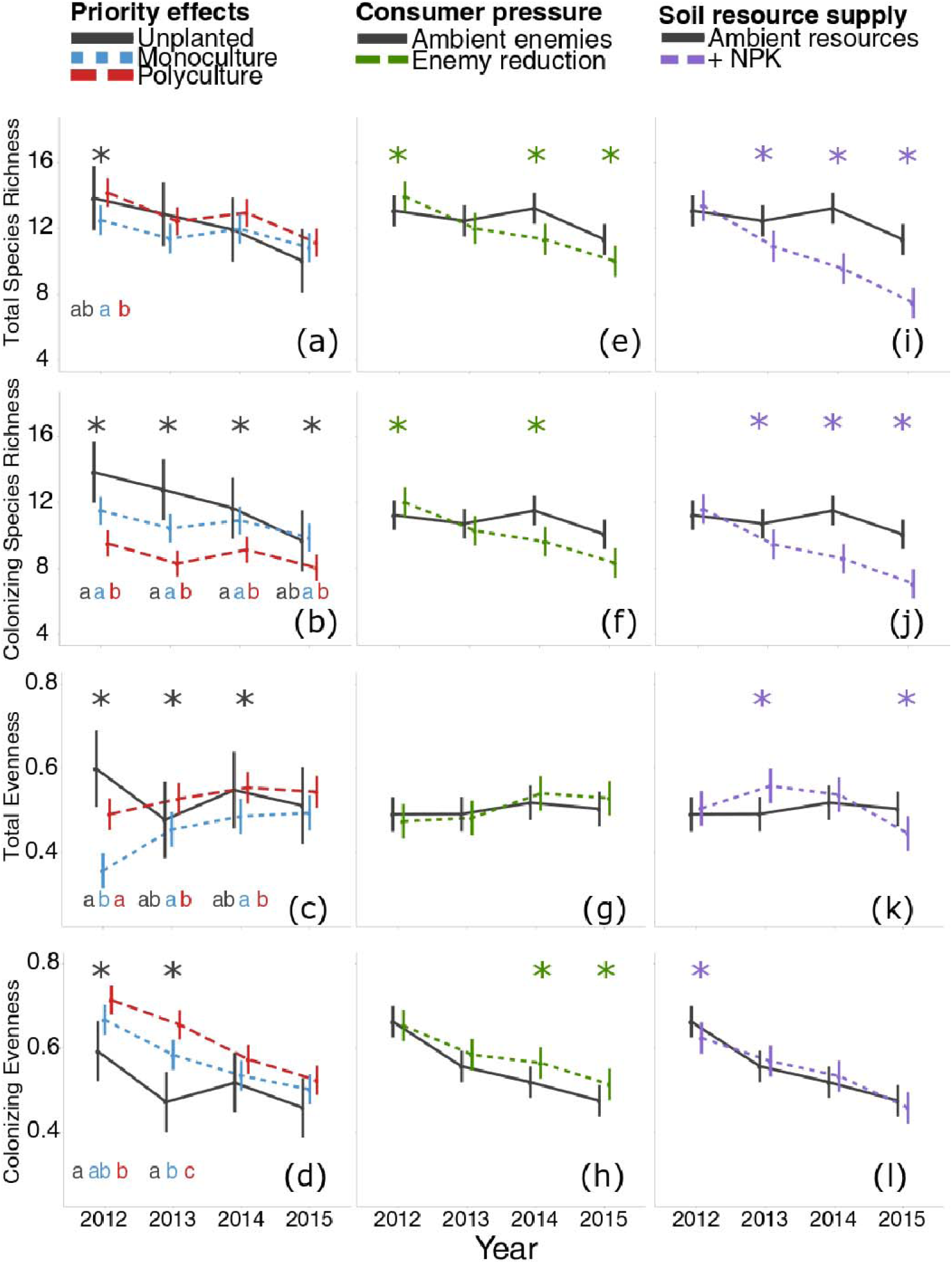
Responses of taxonomic diversity indices to priority effects (initial richness), reduced consumer pressure (pesticide spraying), and increased soil resource supply (fertilization) over four years of succession. Diversity indices are calculated for the entire community (total) and separately for only the non-planted species (colonizing). Error bars are 95% confidence intervals Asterisks indicate significant differences between treatments within years; lower case letters further indicate significant differences between treatment levels for priority effects.

We hypothesized that evenness, or the distribution of resources between species, changes whether species that form early monocultures are able to maintain competitive advantages from a high initial dominance. Consistent with this hypothesis, initial richness influenced total species evenness (p=0.011; Fig. **1c**, Table S5); this effect changed through time (p<0.001; Table S6) and had a significant interaction with fertilization (p<0.01; Table S6). Monocultures had significantly lower evenness than polycultures for the first three years of the study, and lower evenness than unplanted plots in the first year. Colonizing evenness was also influenced by initial richness (p<0.01; Fig **1d**, Table S7), and this effect changed through time (p=0.019; Table S8). The evenness of colonizers was higher in polycultures than monocultures in the first two years of the study. Together, this suggests that species planted as monocultures maintained dominance early in the study, while unplanted plots had similar evenness to polycultures almost immediately. Colonizing species had higher evenness at higher initial richness early in the experiment, suggesting a higher diversity of initial residents limits dominance of any one colonizing species early in succession.

We hypothesized that reducing consumer pressure via spraying would reduce plant diversity during succession as consumers often most strongly limit dominant plant species that may otherwise competitively exclude non-dominant species. Indeed, spraying decreased total species richness in the third and fourth years (Spraying × Time: p=0.011; Fig **1e**, Table S1). Moreover, spraying interacted with fertilization (Spraying × Fertilization: p=0.039, Table S2), and this effect was further modified through time (Spraying × Fertilization × Time: p<0.001; Table S2); spraying only reduced richness in unfertilized plots in later years. This pattern was the same for colonizing species’ richness (Spraying: p<0.01; Spraying × Year: p<0.01; Spraying × Year × Fertilization: p<0.01; Fig **1f**; Table S3, S4). Spraying did not change total species evenness (p=0.49; Fig. **1g**, Table S5). However, spraying did influence the evenness of colonizing species, (p<0.01; Fig. **1h**, Table S7). Despite not having a significant interaction with time (p=0.073; Table S8), this effect was only apparent in the third and fourth years, where colonizing species had higher evenness in sprayed plots. The observed spraying effects on richness are consistent with reduced consumer pressure allowing previously enemy-suppressed species to competitively exclude others, although the temporary increase in colonizing evenness indicates an absence of dominance that would be expected to accompany this.

We hypothesized that increasing soil resource supply via fertilization would lead to increased dominance of one or a few species and competitive exclusion of non-dominants, as resource limitation promotes coexistence. We found that fertilization decreased total species richness (p<0.0001; Fig **1i**, Table S1). This started in the second year and the effect size grew each year (p<0.0001; Table S2). Likewise, fertilization decreased colonizing species richness (p<0.0001; Fig **1j**, Table S3), again starting in the second year and with an increasing effect size through time (p<0.0001; Table S4). Fertilization also influenced total species evenness (p<0.0001; Fig **1k**, Table S5), and this effect changed through time (p<0.0001; Table S6). Here, however, the effect inverted over time, with fertilization increasing total evenness in the second year, showing no effect in the third year, and decreasing total evenness in the fourth year of study. Fertilization also influenced colonizing species evenness through time (p=0.038; Fig 1l; Table S7, S8), but unlike with total evenness, colonizer evenness declined in the first year when fertilized and had no subsequent effects. The strong decreases in richness are consistent with increased soil resources enabling competitive dominance and exclusion by previously resource limited species. The effects on evenness are especially interesting, indicating that colonizing species may have equilibrated in abundance with planted species benefitting from priority effects (i.e. increasing evenness), before overwhelming them in later years and competitively dominating plots in the last year of study.

### Priority effects, consumer pressure, and soil resource supply effects on species resource acquisition and allocation

We hypothesized that increases in community seed mass, reflecting increasing competitive environments for colonizing seedlings, would occur with more diverse priority effects due to higher resource drawdown. Initial richness influenced the CWM of colonizing species’ seed mass (p=0.036; Fig. **2a**, Table S9), and this effect changed through time (p=0.038; Table S10). Specifically, in the first three years of the study, unplanted plots had smaller seeded species colonizing them than polycultures; a significant decrease of seed mass in unplanted plots relative to monocultures was observed only in the first year. Initial richness effects on seed mass were only detectable in unfertilized plots (Fertilization × Initial Richness: p<0.01; Table S10). We additionally hypothesized that resource competition would increase seed mass as competition for belowground resources switched to asymmetric competition for light, requiring more competitive seedlings. Consistent with this hypothesis, fertilization increased colonizer seed mass (p<0.0001, Table S9, Fig **2c**), an effect that was first observed in year two and onward (Fertilization × Time: p<0.0001, Table S10). Spraying did not influence colonizer seed mass (p=0.84, Fig **2b**, Table S9) in any year (p=0.15, Table S10). Together, these results suggest that heavier-seeded species were better able to colonize high richness plots early in the study, while fertilization led to an advantage for heavier-seeded species in later years of the study.

**Fig 2.**
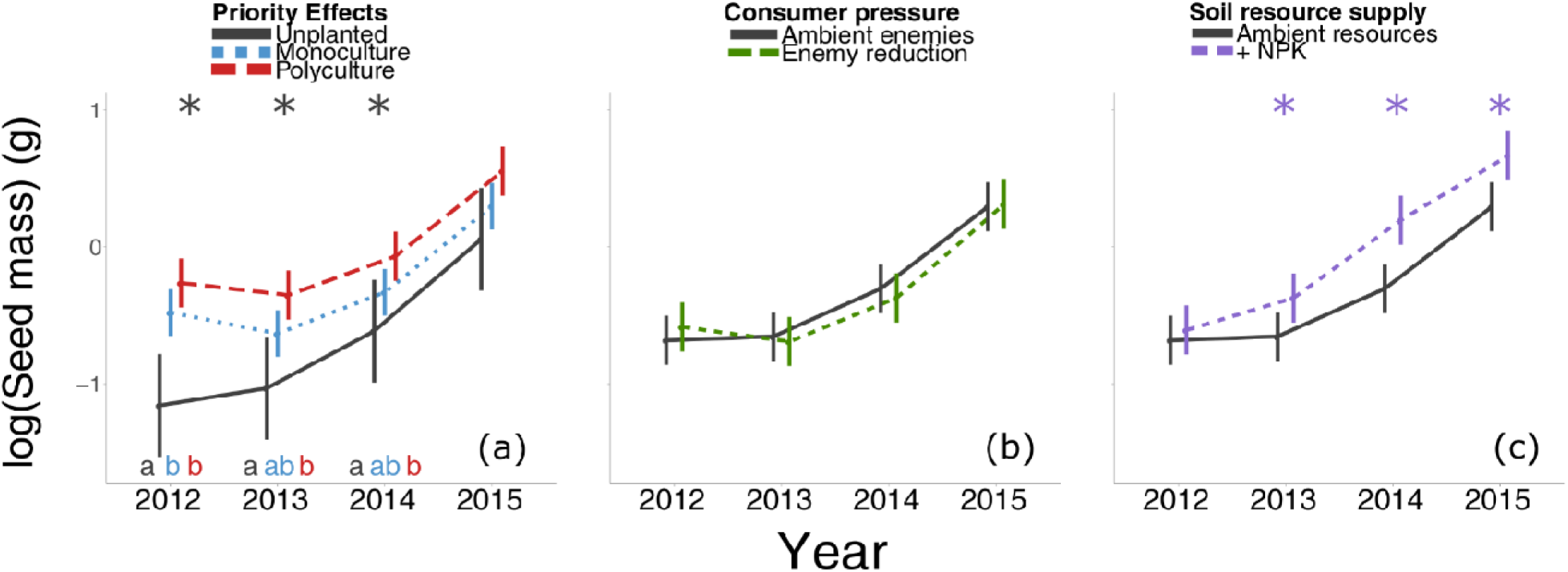
Responses of community weighted mean seed mass of colonizing species to priority effects (initial richness), reduced consumer pressure (pesticide spraying), and increased soil resource supply (fertilization) over four years of succession. Seed mass is a species mean from a database and log-transformed. Error bars are 95% confidence intervals. Asterisks indicate significant differences between treatments within years; lower case letters further indicate significant differences between treatment levels for priority effects.

We hypothesized that increased initial richness, reduced consumer pressure and increased soil resource supply would benefit species with resources acquisitive strategies (i.e. high height and SLA). We additionally tested whether CWMs of traits changed because species were excluded due to poor relative performance (i.e. interspecific) or because species shifted their resource acquisition strategy to match the environment (i.e. intraspecific). In the third year of study when height and SLA were measured, initial richness did not influence interspecific height (p=0.33; Fig **3a**, Table S11), intraspecific height (p=0.30; Fig **3b**, Table S12), interspecific SLA (p=0.47; Fig. **3c**, Table S13), or intraspecific SLA (p=0.30; Fig. **3d**, Table S14). This indicates initial richness did not influence community membership based on these traits, though it is possible these were important traits in the first years of study when initial richness more clearly influenced taxonomic diversity. Spraying did not influence interspecific height (p=0.15; Fig **3a**, Table S11), but did increase intraspecific height (p<0.001; Fig **3b**, Table S12), indicating that spraying did not filter species based on height, but residents were able to grow taller in sprayed plots. Spraying decreased interspecific SLA (p<0.001; Fig **3c**, Table S13), but had no effect on intraspecific SLA (p=0.42; Fig **3d**, Table S14). This is a surprising result contrary to our hypothesis, as it indicates species with thicker, resource conservative leaves were benefitted more by spraying than resource acquisitive leafed species. Fertilization increased both interspecific height (p<0.0001; Fig **3a**, Table S11) and intraspecific height (p<0.0001; Fig **3b**, Table S12). Fertilization increased interspecific SLA (p<0.001; Fig **3c**, Table S13), but had no effect on intraspecific SLA (p=0.089; Fig **3d**, Table S14). This is consistent with the observed diversity effects, suggesting that fertilization gave competitive advantages towards resource acquisitive species with taller stature and more leaf area per unit mass.

**Fig 3.**
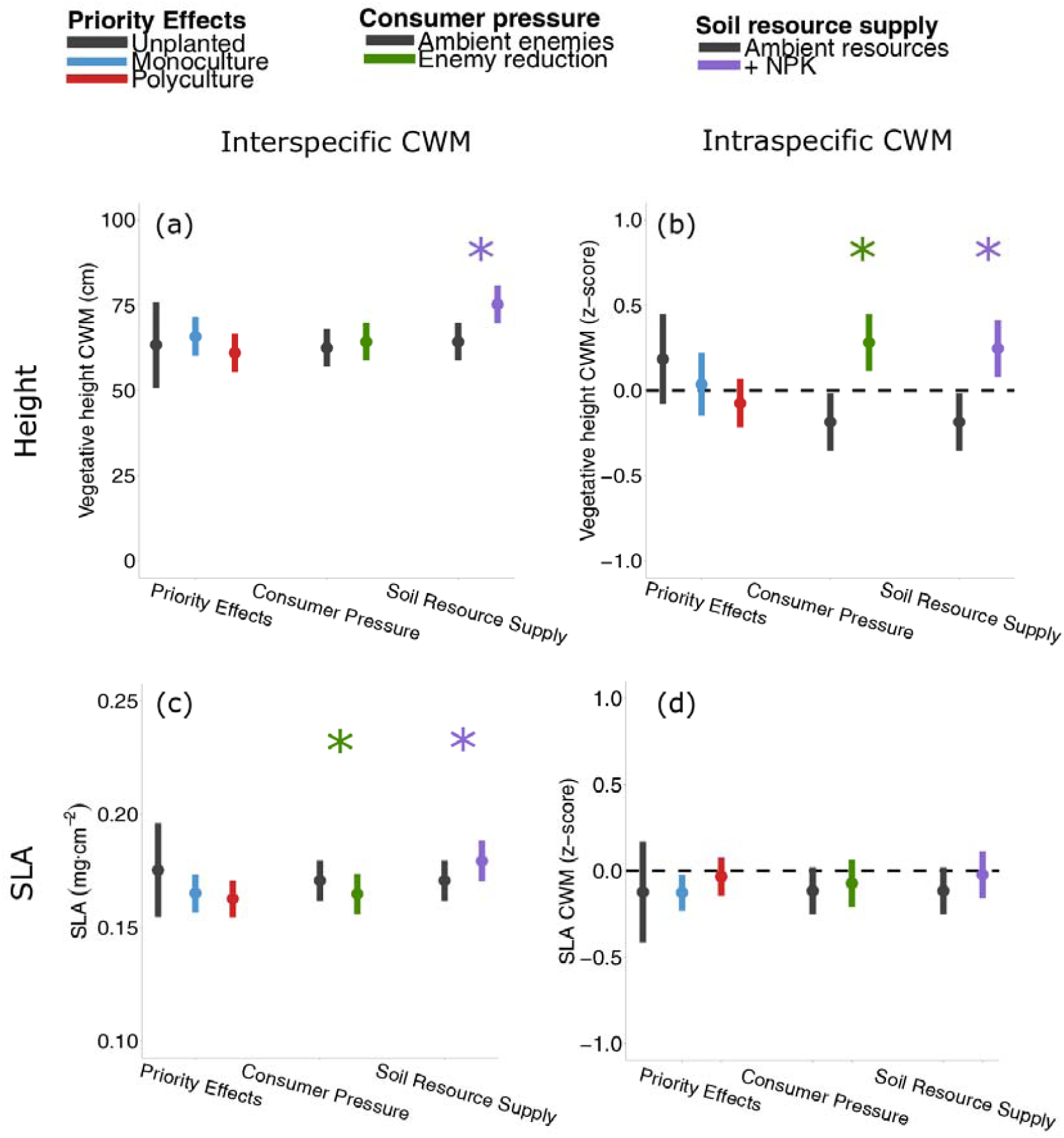
Community height and SLA responses to priority effects (initial richness), reduced consumer pressure (pesticide spraying), and increased soil resource supply (fertilization) after three years of treatment (2014). Trait metrics are split into interspecific community weighted means, calculated using the field measured species-level means, and intraspecific community weighted means, calculated using plot measured species-level z-scores (standardized variation away from field measured species-level means). Error bars are 95% confidence intervals. Asterisks indicate significant differences between treatments levels within treatments.

### Priority effects, consumer pressure, and soil resource supply effects on competition for light

Light attenuation can serve as a proxy for competition for light, which we hypothesized to be a key mechanism associated with diversity changes and our studied drivers of succession. Consistent with this hypothesis, initial richness influenced light attenuation interactively with time (p<0.0001; Fig **4a**; Table S15, S16); in the first year of study, unplanted plots had higher light availability at ground level than monocultures, which had higher light availability than polycultures. No significant differences were observed in subsequent years. Spraying reduced light availability at ground level (p<0.0001, Fig **4b**, Table S15); this occurred in all years but the relative effect size increased in later years (p<0.0001, Table S16). Fertilization decreased light availability at ground level (p<0.0001, Fig **4c**, Table S15), the effect size varied through time (p<0.0001, Table S16), with the greatest decrease in the third year. Spraying and fertilization had a significant interaction (p<0.0001, Table S16), as total reduction of light availability was non-additive (i.e. fertilization did not further decrease light availability in sprayed plots and vice-versa). Light attenuation also explained taxonomic and functional diversity metrics. Light attenuation affected species richness interactively with time (p<0.01), with species richness increasing with light availability from the second year on (Fig. S1). Conversely, evenness only increased with light availability in the first year (Light × Year: p<0.001; Fig. S2). Similarly, CWM seed mass had a negative relationship with light availability only in the first year (Light × Year: p<0.01; Fig S3). In 2014, light availability had a negative relationship with CWM height (p<0.001; Fig S4), but no relationship with CWM SLA (p=0.22; Fig S5). Together, these results indicate that competition for light is an important mechanism, explaining how the treatments we applied ultimately influenced species diversity, height, and seed responses. Higher initial richness, decreased consumer pressure, and increased soil resource supply all decreased light availability in at least the first year. Additionally, these decreases were associated with fewer tall, large-seeded, more dominant species coexisting.

**Fig 4.**
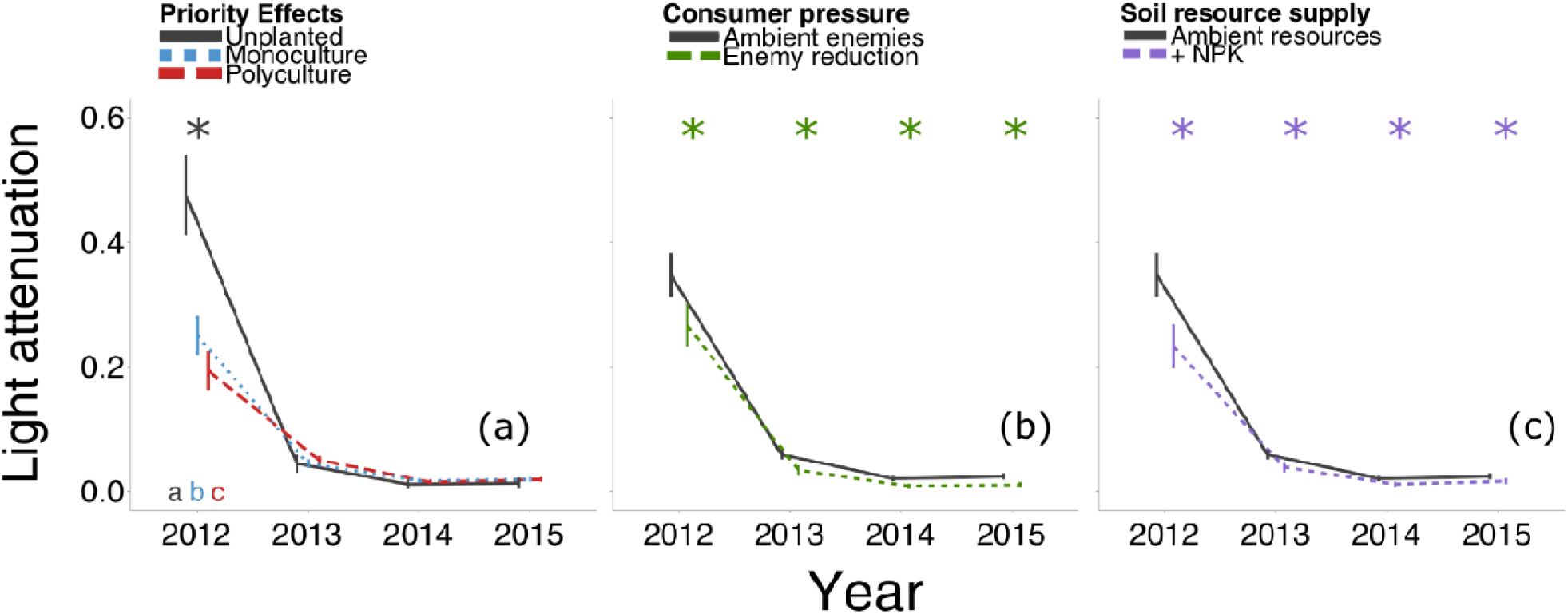
Light availability responses to priority effects (initial richness), reduced consumer pressure (pesticide spraying), and increased soil resource supply (fertilization) over four years of succession. Light availability was measured as attenuation, or the ratio of photosynthetically active radiation above the canopy to the ground. Error bars are 95% confidence intervals. Letters indicate significant differences between treatments within years. Asterisks indicate significant differences between treatments within years; lower case letters further indicate significant differences between treatment levels for priority effects.

## Discussion

Collectively, our results show how priority effects, consumer pressure, and soil resource supply can impact plant secondary succession. Importantly, this experiment reveals that these three drivers are largely additive in effect rather than interactive and can have temporally heterogeneous effects. The drivers generally acted independently from one another, although increased soil resource supply overwhelmed the effects of the other drivers at times. By combining an experimental manipulation, detailed surveys over time, and measures of plant morphological traits, our results show that biotic and abiotic drivers can independently alter successional trajectories by shifting community composition, and that community membership depends on specific traits related to dispersal and resource acquisition and allocation.

The strength of priority effects is modified by the richness of an early successional system and can impact the further development of community diversity at later successional periods (Mouquet *et al.*, 2004). In our experiment, initial richness, which represented a variety of priority effect scenarios including the absence of planted species, only temporarily influenced subsequent richness in the first year, suggesting that the ultimate number of locally co-existing species is governed by other abiotic or biotic processes. Unlike richness, the dominance of planted species in monocultures persisted for 3-years on average, but eventually species evenness also became indistinguishable among differing priority effect scenarios. Although the dominance of planted species decayed over time, the richness of colonizing species was suppressed in polycultures compared to monocultures in the last year of the study, indicating that the maintenance of priority effects of early establishing perennial species in succession was more likely at higher richness. Roscher et al. (2014) similarly found high initial divergence in richness among planted communities following seed addition, followed by convergence of richness in later years, while compositional differences were maintained throughout. Our study shows that this process also occurs with natural colonization. Other studies have similarly found that priority effects of seed arrival times can alter future composition (Körner *et al.*, 2008), even though richness effects are similarly transient (Weidlich *et al.*, 2017). Such persistent composition changes can have lasting impacts on ecosystem functions such as parasite dynamics (Halliday *et al.*, 2019) and biomass production (von Gillhaussen *et al.*, 2014), even when metrics such as richness appear to indicate invariant community diversity.

Defining what diversity precisely *is* has a long history in community ecology (Peet, 1974; Chase *et al.*, 2018), but it remains clear that many dimensions of diversity are important for understanding the role of diversity in ecosystem functions (Naeem *et al.*, 2012). In our study, decomposing taxonomic diversity into richness and evenness revealed competing drivers of succession. On one hand, richness, which indicates how well abiotic and biotic conditions support the co-occurrence of species (Peet, 1974), was depressed when soil resource limitation and consumer pressure were alleviated. Conversely, evenness, an indicator of how equally resources are partitioned within co-occurring species (Peet, 1974), was depressed in response to increased soil resource supply, but remained neutral in response to consumer pressure alleviation, even increasing with respect to colonizing species. Thus, the species richness loss following increased soil resource supply indicates competitive exclusion resulting from one or a few dominant species, while species exclusion from consumer pressure alleviation, unaccompanied by dominance increases, indicated a community-wide suppression by more evenly distributed species. Similarly, decomposing functional diversity into its inter and intraspecific sources of variation revealed otherwise masked responses (e.g. Funk et al., 2017). Examining these components of meaningful functional traits alongside our multifaceted taxonomic diversity results complements our understanding of succession in this system.

Community changes in vegetative height, SLA, and seed mass demonstrated how priority effects, consumer pressure, and resource supply acted on species dispersal and resource acquisition and allocation tradeoffs (Westoby, 1998). Increased soil resource supply led to the strongest effects on trait responses, with clear increases in inter and intraspecific height, interspecific SLA, and seed mass through time. All of these community-level responses to soil resource supply are consistent with increased light competition, as species allocate resources to placing more leaf area at higher parts of the canopy and producing seeds that may be better able to germinate in low light environments (Manning *et al.*, 2009). This is further supported by the observed decrease in light availability. Additionally, increased soil nutrients may favor species with higher specific leaf area (SLA), suggesting a shift towards resource acquisitive strategies (Knops & Reinhart, 2000; Laliberte *et al.*, 2012). We find strong support for this hypothesis, in contrast with several other studies that have been unable to detect this relationship (Wright & Sutton-Grier, 2012; Kazakou *et al.*, 2014).

Unlike increased soil resource supply, reduced consumer pressure decreased interspecific SLA, had no effect on seed mass, and increased intraspecific height without interspecific height shifts. These results are counter-intuitive. Previous work demonstrated that low SLA species are less palatable, and thus less vulnerable to consumer pressure (Schädler *et al.*, 2003)). However, reducing consumer pressure in our experiment actually shifted communities *towards* lower SLA species, suggesting heightened vulnerability to local fungal pathogens and/or insect herbivores. While SLA is correlated with leaf traits conferring consumer resistance (Agrawal & Fishbein, 2006; Moles *et al.*, 2011), its utility as a stand-alone indicator may be limited (Pérez-Harguindeguy *et al.*, 2003; Poorter *et al.*, 2009). Our study indicates that not only did low SLA not provide consumer resistance, but low SLA species actually increased both their height and local abundance when consumer pressure was alleviated. Resource conservative species that invest more into individual leaves depend on increasing leaf longevity to balance the investment (Wright *et al.*, 2004); this along with improved species performance suggests low SLA provided a benefit that was mitigated by consumer pressure.

Increased soil resource supply and reduced consumer pressure both resulted in decreased species richness and light availability, but the differences in interspecific trait shifts indicate they influenced the successional trajectory by acting on different species-level tradeoffs. Vertebrate herbivory suppression was shown to benefit the same species as fertilization in a globally distributed experiment, indicative of a competition-defense tradeoff (Lind *et al.*, 2013). Our results are more consistent with La Pierre et al., (2015), who found that reduced invertebrate herbivore pressure shifted community composition in a different direction from increased resource supply, indicative of a growth-defense tradeoff. Moreover, different invertebrate clades, such as insects and mollusks, may themselves have differing effects on plant composition (Allan & Crawley, 2011), and plant fungal pathogens may filter an entirely different suite of species than animal consumers within the same system (Allan *et al.*, 2010). Similarly, individual soil resources themselves likely have differing effects on species composition (Harpole *et al.*, 2016). Consequently, because priority effects, consumer pressure, and soil resource supply influence a variety of species-level tradeoffs, our results indicate that the multidimensionality of these factors could be an important underlying driver of species coexistence.

Diminishing light availability is often the most evident and rapid change in resource availability during plant community succession (Chase & Leibold, 2003). While fertilization and spraying did not affect community diversity until later years, their effects on light attenuation were immediate. Additionally, increased soil resource supply changed the early relationship between initial richness and light attenuation; in the first year, polycultures drew down light availability more than monocultures, but only in fertilized plots. This suggests that polycultures more rapidly increased light competition than monocultures, but only when soil resource limitations were removed. These observations establish that the lack of an observed community shift, in richness or other community properties, does not indicate an absence of ecosystem responses (Jentsch *et al.*, 2011). In our study, community level shifts lagged a season or more behind observed changes in light availability responding to changes in soil resource supply and consumer pressure, suggesting changes in ecosystem responses may help detect important community level responses before they occur.

Our results showed consumer pressure effects being suppressed by increased soil resource supply, contrasting previous experimental work in another herbaceous system. Specifically, our results indicate that soil resources were more limiting than consumers, whereas Allan & Crawley, (2011) found that consumers were more limiting than soil resources. These apparently contrasting results might occur because neither study was designed to measure multiple treatment levels and the shape of the nonlinearity in the nutrient-enemy interaction that might result. Furthermore, to some extent the outcome of ecological processes will always be contingent on system characteristics (Gruner *et al.*, 2008). Valuable next steps in exploring interaction landscapes (i.e. at what intensity do drivers interact and under what conditions) include shifting towards experiments emphasizing gradients of treatment levels as opposed to one or a few treatment levels at high replication (Kreyling *et al.*, 2018) and continued replication of experiments across geographic scales (Borer *et al.*, 2014a). In any case, system contingencies provide an opportunity to better understand ecological processes like succession.

An advantage to studying successional trajectories in the Southeastern US is that this system undergoes succession more rapidly than more northern climates, due to reduced temperature constraints (Fridley & Wright, 2018). Consequently, this study, which took place over only four years, represented the full transition from bare soil through early herbaceous dominance and into the beginning of woody encroachment. Thus, by focusing our experiment on a single field system, we were able to address a question that would be intractable or impossible in many other systems, where succession proceeds more slowly.

Our four year study revealed that priority effects, consumer pressure, and soil resource supply largely influence the trajectory of succession independently from one another, acting at different points in time, on different components of diversity, and on different species traits. Understanding succession is therefore likely to be incomplete when single drivers, time points, or diversity responses are taken under consideration. Moreover, changes in light availability preceded changes in diversity in response to increased soil resource supply and decreased consumer pressure by a full year, indicating a lag effect to a more immediate shift in resource limitation. Collectively, our results indicate that abiotic and biotic controls of successional trajectories are temporally heterogeneous and can drive succession in different compositional directions by favoring species with specific resource acquisition, defense, and dispersal strategies.

## Acknowledgements

We thank Charles Mitchell for considerable support and guidance on designing and implementing the study. Justin Wright, John Bruno, and Peter White provided helpful advice and discussion. Many members of the Plant Ecology Lab, Mitchell Lab, and Duke Forest provided field assistance and botanical expertise. PAW, FWH, and RWH were supported by UNC’s Alma Holland Beers Scholarship and WC Coker Fellowship. RWH and FWH were supported by the UNC Graduate School Dissertation Completion Fellowship. FWH was supported by the NSF Graduate Research Fellowship.

## Author contributions

PAW, FWH, and RWH conceived and implemented the experiment. PAW analyzed the data and wrote the first draft. All authors contributed to revisions of the manuscript.

## References

Agrawal AA, Fishbein M. 2006. Plant defense syndromes. Ecology 87: S132–S149.

Allan E, Crawley MJ. 2011. Contrasting effects of insect and molluscan herbivores on plant diversity in a long-term field experiment. Ecology letters 14: 1246–53.

Allan E, van Ruijven J, Crawley MJ. 2010. Foliar fungal pathogens and grassland biodiversity. Ecology 91: 2572–82.

Borer ET, Harpole WS, Adler PB, Lind EM, Orrock JL, Seabloom EW, Smith MD. 2014a. Finding generality in ecology: a model for globally distributed experiments (R Freckleton, Ed.). Methods in Ecology and Evolution 5: 65–73.

Borer ET, Seabloom EW, Gruner DS, Harpole WS, Hillebrand H, Lind EM, Adler PB, Alberti J, Anderson TM, Bakker JD, et al. 2014b. Herbivores and nutrients control grassland plant diversity via light limitation. Nature 508: 517–20.

Chase JM, Leibold MA. 2003. Ecological niches□: linking classical and contemporary approaches.

Chase JM, McGill BJ, McGlinn DJ, May F, Blowes SA, Xiao X, Knight TM, Purschke O, Gotelli NJ. 2018. Embracing scale-dependence to achieve a deeper understanding of biodiversity and its change across communities. Ecology Letters 21: 1737–1751.

Coley PD, Bryant JP, Chapin FS. 1985. Resource Availability and Plant Antiherbivore Defense. Science 230: 895–899.

DeMalach N, Zaady E, Kadmon R. 2017. Light asymmetry explains the effect of nutrient enrichment on grassland diversity. Ecology Letters 20: 60–69.

Dickson T, Foster B. 2011. Fertilization decreases plant biodiversity even when light is not limiting. Ecology letters 14: 380–8.

Dickson T, Mittelbach G, Reynolds H, Gross K. 2014. Height and clonality traits determine plant community responses to fertilization. Ecology 95: 2443–2452.

Fargione JE, Tilman D. 2005. Diversity decreases invasion via both sampling and complementarity effects. Ecology Letters 8: 604–611.

Fridley JD, Wright JP. 2018. Temperature accelerates the rate fields become forests. Proceedings of the National Academy of Sciences of the United States of America 115: 4702–4706.

Funk JL, Larson JE, Ames GM, Butterfield BJ, Cavender □Bares J, Firn J, Laughlin DC, Sutton □Grier AE, Williams L, Wright JP. 2017. Revisiting the H oly G rail: using plant functional traits to understand ecological processes. Biological Reviews 92: 1156–1173.

von Gillhaussen P, Rascher U, Jablonowski ND, Plückers C, Beierkuhnlein C, Temperton VM. 2014. Priority Effects of Time of Arrival of Plant Functional Groups Override Sowing Interval or Density Effects: A Grassland Experiment (M Heil, Ed.). PLoS ONE 9: e86906.

Grime J. 1973. Competitive exclusion in herbaceous vegetation. Nature 242: 344–347.

Gruner DS, Smith JE, Seabloom EW, Sandin SA, Ngai JT, Hillebrand H, Harpole WS, Elser JJ, Cleland EE, Bracken MES, et al. 2008. A cross-system synthesis of consumer and nutrient resource control on producer biomass. Ecology Letters 11: 740–755.

Halliday FW, Heckman RW, Wilfahrt PA, Mitchell CE. 2017. A multivariate test of disease risk reveals conditions leading to disease amplification. Proceedings of the Royal Society B: Biological Sciences 284.

Halliday FW, Heckman RW, Wilfahrt PA, Mitchell CE. 2019. Past is prologue: host community assembly and the risk of infectious disease over time. Ecology Letters 22: 138–148.

Harpole WS, Sullivan LL, Lind EM, Firn J, Adler PB, Borer ET, Chase J, Fay PA, Hautier Y, Hillebrand H, et al. 2016. Addition of multiple limiting resources reduces grassland diversity. Nature 537: 93–96.

Harpole WS, Tilman D. 2007. Grassland species loss resulting from reduced niche dimension. Nature 446: 791–793.

Heckman RW, Halliday FW, Wilfahrt PA, Mitchell CE. 2017. Effects of native diversity, soil nutrients, and natural enemies on exotic invasion in experimental plant communities. Ecology 98: 1409–1418.

Heckman RW, Wright JP, Mitchell CE. 2016. Joint effects of nutrient addition and enemy exclusion on exotic plant success. Ecology 97: 3337–3345.

Hodapp D, Borer ET, Harpole WS, Lind EM, Seabloom EW, Adler PB, Alberti J, Arnillas CA, Bakker JD, Biederman L, et al. 2018. Spatial heterogeneity in species composition constrains plant community responses to herbivory and fertilisation. Ecology Letters 21: 1364–1371.

Hooper DU, Vitousek PM. 1998. Effects of plant composition and diversity on nutrient cycling. Ecological Monographs 68: 121–149.

Isbell F, Reich PB, Tilman D, Hobbie SE, Polasky S, Binder S. 2013. Nutrient enrichment, biodiversity loss, and consequent declines in ecosystem productivity. Proceedings of the National Academy of Sciences.

Jentsch A, Kreyling J, Elmer M, Gellesch E, Glaser B, Grant K, Hein R, Lara M, Mirzae H, Nadler SE, et al. 2011. Climate extremes initiate ecosystem-regulating functions while maintaining productivity. Journal of Ecology 99: 689–702.

Kazakou E, Violle C, Roumet C, Navas M-L, Vile D, Kattge J, Garnier E. 2014. Are trait-based species rankings consistent across data sets and spatial scales? Journal of Vegetation Science 25: 235–247.

Kempel A, Razanajatovo M, Stein C, Unsicker SB, Auge H, Weisser WW, Fischer M, Prati D. 2015. Herbivore preference drives plant community composition. Ecology 96: 2923–2934.

Knops JMH, Reinhart K. 2000. Specific Leaf Area Along a Nitrogen Fertilization Gradient. The American Midland Naturalist 144: 265–272.

Koerner SE, Smith MD, Burkepile DE, Hanan NP, Avolio ML, Collins SL, Knapp AK, Lemoine NP, Forrestel EJ, Eby S, et al. 2018. Change in dominance determines herbivore effects on plant biodiversity. Nature Ecology & Evolution 2: 1925–1932.

Körner C, Stöcklin J, Reuther-Thiébaud L, Pelaez-Riedl S. 2008. Small differences in arrival time influence composition and productivity of plant communities. New Phytologist 177: 698–705.

Kreyling J, Schweiger AH, Bahn M, Ineson P, Migliavacca M, Morel-Journel T, Christiansen JR, Schtickzelle N, Larsen KS. 2018. To replicate, or not to replicate - that is the question: how to tackle nonlinear responses in ecological experiments. Ecology Letters 21: 1629–1638.

Laliberte E, Shipley B, Norton DA, Scott D. 2012. Which plant traits determine abundance under long-term shifts in soil resource availability and grazing intensityl_J ? Journal of Ecology 100: 662–677.

Leishman MR, Wright I, Moles AT, Westoby M. 2000. The evolutionary ecology of seed size. In: Fenner M, ed. Seeds: The Ecology of Regeneration in Plant Communities. New York, NY: CAB International, 31–58.

Lenth R, Hervé M. 2015. lsmeans: Least-Squares Means. R package version 2.16.

Levine J, D’Antonio C. 1999. Elton revisited: a review of evidence linking diversity and invasibility. Oikos 87: 15–26.

Lind EM, Borer E, Seabloom E, Adler P, Bakker JD, Blumenthal DM, Crawley M, Davies K, Firn J, Gruner DS, et al. 2013. Life-history constraints in grassland plant species: a growth-defence trade-off is the norm. Ecology letters 16: 513–21.

Manning P, Houston K, Evans T. 2009. Shifts in seed size across experimental nitrogen enrichment and plant density gradients. Basic and Applied Ecology 10: 300–308.

Moles AT, Wallis IR, Foley WJ, Warton DI, Stegen JC, Bisigato AJ, Cella-Pizarro L, Clark CJ, Cohen PS, Cornwell WK, et al. 2011. Putting plant resistance traits on the map: a test of the idea that plants are better defended at lower latitudes. New Phytologist 191: 777–788.

Mordecai EA. 2011. Pathogen impacts on plant communities: unifying theory, concepts, and empirical work. Ecological Monographs 81: 429–441.

Mortensen B, Danielson B, Harpole WS, Alberti J, Arnillas CA, Biederman L, Borer ET, Cadotte MW, Dwyer JM, Hagenah N, et al. 2018. Herbivores safeguard plant diversity by reducing variability in dominance (M Heard, Ed.). Journal of Ecology 106: 101–112.

Mouquet N, Leadley P, Mériguet J, Loreau M. 2004. Immigration and local competition in herbaceous plant communities: a three-year seed-sowing experiment. Oikos 1: 77–90.

Naeem S, Duffy JE, Zavaleta E. 2012. The functions of biological diversity in an age of extinction. Science 336: 1401–1406.

Oksanen J, Blanchet FG, Kindt R, Legendre P, Minchin PR, O’hara RB, Simpson GL, Solymos P, Stevens MHH, Wagner H. 2013. Package ‘vegan’. Community ecology package, version 2.

Olff H, Ritchie M. 1998. Effects of herbivores on grassland plant diversity. Trends in Ecology & Evolution 13: 261–265.

Peet RK. 1974. The measurement of species diversity. Annual review of ecology and systematics 5: 285–307.

Pérez-Harguindeguy N, Díaz S, Garnier E, Lavorel S, Poorter H, Jaureguiberry P, Bret-Harte M, Cornwell W, Craine JM, Gurvich DE, et al. 2013. New handbook for standardised measurement of plant functional traits worldwide. Australian Journal of Botany.

Pérez-Harguindeguy N, Díaz S, Vendramini F, Cornelissen JHC, Gurvich DE, Cabido M. 2003. Leaf traits and herbivore selection in the field and in cafeteria experiments. Austral Ecology 28: 642–650.

La Pierre KJ, Joern A, Smith MD. 2015. Invertebrate, not small vertebrate, herbivory interacts with nutrient availability to impact tallgrass prairie community composition and forb biomass. Oikos 124: 842–850.

Poorter H, Niinemets Ü, Poorter L, Wright IJ, Villar R. 2009. Causes and consequences of variation in leaf mass per area (LMA): a meta-analysis. New Phytologist 182: 565–588.

Purschke O, Schmid BC, Sykes MT, Poschlod P, Michalski SG, Durka W, Kühn I, Winter M, Prentice HC. 2013. Contrasting changes in taxonomic, phylogenetic and functional diversity during a long-term succession: insights into assembly processes. Journal of Ecology 101: 857–866.

R Core Team. 2015. R: A Language and Environment for Statistical Computing. : https://www.R-project.org/.

Rajaniemi TK, Allison VJ, Goldberg DE. 2003. Root competition can cause a decline in diversity with increased productivity. Journal of Ecology 91: 407–416.

Roscher C, Schumacher J, Gerighausen U, Schmid B. 2014. Different assembly processes drive shifts in species and functional composition in experimental grasslands varying in sown diversity and community history. PLoS ONE 9: e101928.

Royal Botanic Gardens Kew. 2016. Seed Information Database (SID). Version 7.1.

Schädler M, Jung G, Auge H, Brandl R. 2003. Palatability, decomposition and insect herbivory: patterns in a successional old-field plant community. Oikos 103: 121–132.

Schmid B, Hector A, Huston MA, Inchausti P, Nijs I, Leadley PW, Tilman D. 2002. The design and analysis of biodiversity experiments. Biodiversity and ecosystem functioning: synthesis and perspectives: 61–75.

Shipley B, Meziane D. 2002. The balanced-growth hypothesis and the allometry of leaf and root biomass allocation. Functional Ecology 16: 326–331.

Suding KN, Collins SL, Gough L, Clark C, Cleland EE, Gross KL, Milchunas DG, Pennings S. 2005. Functional-and abundance-based mechanisms explain diversity loss due to N fertilization. Proceedings of the National Academy of Sciences of the United States of America 102: 4387–92.

Tilman D. 2004. Niche tradeoffs, neutrality, and community structure: a stochastic theory of resource competition, invasion, and community assembly. Proceedings of the National Academy of Sciences of the United States of America 101: 10854–61.

Tuomisto H. 2012. An updated consumer’s guide to evenness and related indices. Oikos 121: 1203–1218.

Turnbull L, Rees M, Crawley M. 1999. Seed mass and the competition/colonization trade-off: a sowing experiment. Journal of Ecology 87: 899–912.

Veen GF, van der Putten WH, Bezemer TM. 2018. Biodiversity-ecosystem functioning relationships in a long-term non-weeded field experiment. Ecology.

Violle C, Castro H, Richarte J, Navas M-L. 2009. Intraspecific seed trait variations and competition: passive or adaptive response? Functional Ecology 23: 612–620.

Webb CT, Hoeting JA, Ames GM, Pyne MI, Poff N. 2010. A structured and dynamic framework to advance traits-based theory and prediction in ecology. Ecology letters 13: 267–83.

Weidlich EWA, von Gillhaussen P, Delory BM, Blossfeld S, Poorter H, Temperton VM. 2017. The Importance of Being First: Exploring Priority and Diversity Effects in a Grassland Field Experiment. Frontiers in Plant Science 7: 2008.

Weisser WW, Roscher C, Meyer ST, Ebeling A, Luo G, Allan E, Beßler H, Barnard RL, Buchmann N, Buscot F, et al. 2017. Biodiversity effects on ecosystem functioning in a 15-year grassland experiment: Patterns, mechanisms, and open questions. Basic and Applied Ecology 23: 1–73.

Westoby M. 1998. A leaf-height-seed (LHS) plant ecology strategy scheme. Plant and soil 199: 213–227.

Westoby M, Falster DS, Moles AT, Vesk PA, Wright IJ. 2002. Plant ecological strategies: Some leading dimensions of variation between species. Annual Review of Ecology and Systematics 33: 125–159.

Wright IJ, Reich PB, Westoby M, Ackerly DD, Baruch Z, Bongers F, Cavender-Bares J, Chapin T, Cornelissen JHC, Diemer M, et al. 2004. The worldwide leaf economics spectrum. Nature 428: 821–7.

Wright JP, Sutton-Grier A. 2012. Does the leaf economic spectrum hold within local species pools across varying environmental conditions? Functional Ecology 26: 1390–1398.

Züst T, Agrawal AA. 2017. Trade-Offs Between Plant Growth and Defense Against Insect Herbivory: An Emerging Mechanistic Synthesis. Annual Review of Plant Biology 68: 513–534.

Zuur AF, Ieno EN, Walker N, Saveliev AA, Smith GM. 2009. Mixed effects models and extensions in ecology with R. New York, NY: Springer New York.

