## Supplementary material for "Priority effects, consumer pressure, and soil resources independently alter plant diversity and resource strategies during a multi-year successional field experiment"

The following Supporting Information is available for this article:

**Methods S1** Extended methods on planting and spraying treatments, and vegetation surveys.

**Table S1** Model of total species richness

**Table S2** Multiple comparisons of total species richness

**Table S3** Model of colonizing species richness

**Table S4** Multiple comparisons of colonizing species richness

**Table S5** Model of total species evenness

**Table S6** Multiple comparisons of total species evenness

**Table S7** Model of colonizing species evenness

**Table S8** Multiple comparisons of colonizing species evenness

**Table S9** Model of colonizing species seed mass

**Table S10** Multiple comparisons of colonizing species seed mass

**Table S11** Model of interspecific height

**Table S12** Model of intraspecific height

**Table S13** Model of interspecific SLA

**Table S14** Model of intraspecific SLA

**Table S15** Model of light attenuation

**Table S16** Multiple comparisons of light attenuation

**Figure S1** Relationship between total species richness and light attenuation across time.

**Figure S2** Relationship between total species evenness and light attenuation across time.

**Figure S3** Relationship between species' seed mass and light attenuation across time.

**Figure S4** Relationship between species' height and light attenuation in the third year of study.

**Figure S5** Relationship between species' SLA and light attenuation in the third year of study.

**Methods S1**

The planted species pool were six perennial, native, herbaceous species already occurring at Widener Farm: three grasses, *Andropogon virginicus*, *Setaria parviflora*, and *Tridens flavus*; two asters, *Packera anonyma* and *Solidago pinetorum*; and one mint, *Scutellaria integrifolia*. Eight to twelve weeks after planting in the greenhouse, species were transplanted into the field by cutting small holes into the landscape fabric, digging small holes, and planting 41 individuals per plot in a checkerboard fashion. In polycultures, four species were randomly assigned to eight spaces, with a fifth randomly selected species per plot being assigned to nine spaces. *Setaria parviflora* was planted entirely in 2012 to replace a species that experienced 100% mortality in 2011.

The pesticides applied are commonly used in ecological field studies: a foliar fungicide (mancozeb, Dithane® DF, Dow AgroSciences, Indianapolis, IN) and an insecticide (es-fenvalerate, Asana® XL, Dupont, Wilmington, DE).

Vegetation surveys entailed all three authors searching within the subplot area for all rooted vascular plants and any vines present in the subplot, whether or not they were rooted in the plot, before jointly estimating the total percent cover of each species. Plots usually exceeded 100% cover due to canopy overlap between species.

**Table S1** *Total richness model. Summary of most parsimonious interaction model of total richness in response to priority effects, consumer pressure, and soil resource supply across time.*

| *Effect* | *DF (Num,Den)* | *F value* | *P value* |
|---|---|---|---|
| *(Intercept)* | 1,762 | 2680.92 | < 0.001 |
| *Priority* | 2,10 | 2.39 | 0.142 |
| *Consumers* | 1,236 | 1.7 | 0.193 |
| *Soil Resources* | 1,236 | 91.51 | < 0.001 |
| *Year* | 3,762 | 134.48 | < 0.001 |
| *Block* | 4,236 | 44.16 | < 0.001 |
| *Priority:Consumers* | 2,236 | 1.38 | 0.254 |
| *Priority:Soil Resources* | 2,236 | 0.66 | 0.515 |
| *Consumers:Soil Resources* | 1,236 | 4.16 | 0.043 |
| *Priority:Year* | 6,762 | 3.2 | 0.004 |
| *Consumers:Year* | 3,762 | 3.75 | 0.011 |
| *Soil Resources:Year* | 3,762 | 17.72 | < 0.001 |
| *Consumers:Soil Resources:Year* | 3,762 | 6.26 | < 0.001 |

**Table S2** Total richness multiple comparisons. Results of Least-Square Means post-hoc Tukey tests and multiple comparisons on total species richness. The comparison shows the treatment vs. control, in the case of the priority effect treatment this compares the polyculture (P), monoculture (M), and unplanted (U). Interaction indicates whether the effect was in unfertilized plots or fertilized for priority effect and spraying treatments, and unsprayed or sprayed the fertilization treatment. The estimated control is the model estimated value. For priority effect this shows the estimated mean value of the lower planted richness level. Estimated effect size is the estimated effect divided by estimated control for comparisons with a significant value ($p < 0.05$).

| Comparison | Interaction | Year | Estimated Effect | Standard Error | T-ratio | p-value | Estimated Control* | Estimated effect size |
|---|---|---|---|---|---|---|---|---|
| | | | | **Priority Effects** | | | | |
| M vs U | UNFERTILIZED | 2012 | -1.7 | 1.01 | -1.68 | 0.259 | 14.1 | - |
| P vs U | UNFERTILIZED | 2012 | 0.11 | 1 | 0.109 | 0.993 | 14.1 | - |
| P vs. M | UNFERTILIZED | 2012 | 1.81 | 0.548 | 3.31 | 0.020 | 12.4 | 14.6 |
| M vs U | UNFERTILIZED | 2013 | -1.85 | 1.01 | -1.83 | 0.210 | 13.1 | - |
| P vs U | UNFERTILIZED | 2013 | -0.657 | 1 | -0.655 | 0.794 | 13.1 | - |
| P vs. M | UNFERTILIZED | 2013 | 1.2 | 0.548 | 2.18 | 0.123 | 11.3 | - |
| M vs U | UNFERTILIZED | 2014 | -0.319 | 1.01 | -0.315 | 0.947 | 12.2 | - |
| P vs U | UNFERTILIZED | 2014 | 0.793 | 1 | 0.79 | 0.717 | 12.2 | - |
| P vs. M | UNFERTILIZED | 2014 | 1.11 | 0.548 | 2.03 | 0.155 | 11.9 | - |
| M vs U | UNFERTILIZED | 2015 | 0.439 | 1.01 | 0.433 | 0.903 | 10.3 | - |
| P vs U | UNFERTILIZED | 2015 | 0.885 | 1 | 0.882 | 0.663 | 10.3 | - |
| P vs. M | UNFERTILIZED | 2015 | 0.446 | 0.548 | 0.814 | 0.703 | 10.7 | - |
| M vs U | FERTILIZED | 2012 | -0.964 | 1.01 | -0.952 | 0.622 | 13.4 | - |
| P vs U | FERTILIZED | 2012 | 0.557 | 1 | 0.555 | 0.846 | 13.4 | - |
| P vs. M | FERTILIZED | 2012 | 1.52 | 0.548 | 2.78 | 0.047 | 12.5 | 12.2 |
| M vs U | FERTILIZED | 2013 | -1.11 | 1.01 | -1.1 | 0.536 | 11.1 | - |
| P vs U | FERTILIZED | 2013 | -0.21 | 1 | -0.209 | 0.976 | 11.1 | - |
| P vs. M | FERTILIZED | 2013 | 0.904 | 0.548 | 1.65 | 0.270 | 9.96 | - |
| M vs U | FERTILIZED | 2014 | 0.419 | 1.01 | 0.414 | 0.911 | 9.02 | - |
| P vs U | FERTILIZED | 2014 | 1.24 | 1 | 1.24 | 0.460 | 9.02 | - |
| P vs. M | FERTILIZED | 2014 | 0.821 | 0.548 | 1.5 | 0.332 | 9.44 | - |
| M vs U | FERTILIZED | 2015 | 1.18 | 1.01 | 1.16 | 0.500 | 6.91 | - |
| P vs U | FERTILIZED | 2015 | 1.33 | 1 | 1.33 | 0.413 | 6.91 | - |
| P vs. M | FERTILIZED | 2015 | 0.154 | 0.548 | 0.282 | 0.957 | 8.09 | - |
| | | | | | **Consumer pressure** | | | |
| Spraying | UNFERTILIZED | 2012 | 0.626 | 0.44 | 1.42 | 0.157 | 13.2 | - |
| Spraying | UNFERTILIZED | 2013 | -0.708 | 0.44 | -1.61 | 0.11 | 12.6 | - |
| Spraying | UNFERTILIZED | 2014 | -2.12 | 0.44 | -4.81 | 0.000 | 13.4 | -15.8 |
| Spraying | UNFERTILIZED | 2015 | -1.56 | 0.44 | -3.54 | 0.000 | 11.5 | -13.6 |
| Spraying | FERTILIZED | 2012 | -0.317 | 0.44 | -0.72 | 0.472 | 13.9 | - |
| Spraying | FERTILIZED | 2013 | -0.659 | 0.44 | -1.49 | 0.136 | 11.9 | - |
| Spraying | FERTILIZED | 2014 | -0.068 | 0.44 | -0.154 | 0.877 | 11.3 | - |
| Spraying | FERTILIZED | 2015 | 0.433 | 0.44 | 0.983 | 0.326 | 9.95 | - |
| | | | | | **Soil resource supply** | | | |
| Fertilization | UNSPRAYED | 2012 | 0.199 | 0.44 | 0.452 | 0.651 | 13.2 | - |
| Fertilization | UNSPRAYED | 2013 | -1.67 | 0.44 | -3.79 | 0.000 | 12.6 | -13.2 |
| Fertilization | UNSPRAYED | 2014 | -3.79 | 0.44 | -8.59 | 0.000 | 13.4 | -28.3 |
| Fertilization | UNSPRAYED | 2015 | -3.97 | 0.44 | -9.02 | 0.000 | 11.5 | -34.5 |
| Fertilization | SPRAYED | 2012 | -0.744 | 0.44 | -1.69 | 0.093 | 13.9 | - |
| Fertilization | SPRAYED | 2013 | -1.62 | 0.44 | -3.68 | 0.000 | 11.9 | -13.6 |
| Fertilization | SPRAYED | 2014 | -1.73 | 0.44 | -3.93 | 0.000 | 11.3 | -15.4 |
| Fertilization | SPRAYED | 2015 | -1.98 | 0.44 | -4.5 | 0.000 | 9.95 | -19.9 |

**Table S3** Colonizing richness model. Summary of most parsimonious interaction model of colonizing richness in response to priority effects, consumer pressure, and soil resource supply across time.

| *Effect* | *DF (Num,Den)* | *F value* | *P value* |
|---|---|---|---|
| *(Intercept)* | 1,762 | 2437.34 | < 0.001 |
| *Priority* | 2,10 | 2.38 | 0.143 |
| *Consumers* | 1,236 | 6.79 | 0.01 |
| *Soil Resources* | 1,236 | 44 | < 0.001 |
| *Year* | 3,762 | 75.41 | < 0.001 |
| *Block* | 4,236 | 47.52 | < 0.001 |
| *Priority:Consumers* | 2,236 | 1.4 | 0.249 |
| *Priority:Soil Resources* | 2,236 | 0.44 | 0.647 |
| *Consumers:Soil Resources* | 1,236 | 1.15 | 0.284 |
| *Priority:Year* | 6,762 | 3.09 | 0.005 |
| *Consumers:Year* | 3,762 | 3.67 | 0.012 |
| *Soil Resources:Year* | 3,762 | 7.89 | < 0.001 |
| *Consumers:Soil Resources:Year* | 3,762 | 5.85 | 0.001 |

**Table S4** Colonizing richness multiple comparisons. Results of Least-Square Means post-hoc Tukey tests and multiple comparisons on colonizing species richness. The comparison shows the treatment vs. control, in the case of the priority effect treatment this compares the polyculture (P), monoculture (M), and unplanted (U). Interaction indicates whether the effect was in unfertilized plots or fertilized for priority effect and spraying treatments, and unsprayed or sprayed for the fertilization treatment. The estimated control is the model estimated value. For priority effect this shows the estimated mean value of the lower planted richness level. Estimated effect size is the estimated effect divided by estimated control for comparisons with a significant value ($p < 0.05$).

| Comparison | Interaction | Year | Estimated Effect | Standard Error | DF | T-ratio | p-value | Estimated Control* | Estimated effect size |
|---|---|---|---|---|---|---|---|---|---|
| **Priority Effects** | | | | | | | | | |
| M vs U | UNFERTILIZED | 2012 | -2.75 | 0.97 | 10 | -2.83 | 0.043 | 14.3 | -19.3 |
| P vs U | UNFERTILIZED | 2012 | -4.84 | 0.956 | 10 | -5.06 | 0.001 | 14.3 | -33.9 |
| P vs. M | UNFERTILIZED | 2012 | -2.09 | 0.518 | 10 | -4.03 | 0.006 | 11.5 | -18.1 |
| M vs U | UNFERTILIZED | 2013 | -2.76 | 0.97 | 10 | -2.84 | 0.042 | 13.2 | -20.9 |
| P vs U | UNFERTILIZED | 2013 | -5 | 0.956 | 10 | -5.23 | 0.001 | 13.2 | -37.8 |
| P vs. M | UNFERTILIZED | 2013 | -2.24 | 0.518 | 10 | -4.32 | 0.004 | 10.4 | -21.4 |
| M vs U | UNFERTILIZED | 2014 | -1.17 | 0.97 | 10 | -1.2 | 0.479 | 12.1 | - |
| P vs U | UNFERTILIZED | 2014 | -3.05 | 0.956 | 10 | -3.19 | 0.024 | 12.1 | -25.2 |
| P vs. M | UNFERTILIZED | 2014 | -1.88 | 0.518 | 10 | -3.63 | 0.012 | 10.9 | -17.2 |
| M vs U | UNFERTILIZED | 2015 | -0.232 | 0.97 | 10 | -0.24 | 0.969 | 10.1 | - |
| P vs U | UNFERTILIZED | 2015 | -2.15 | 0.956 | 10 | -2.25 | 0.110 | 10.1 | - |
| P vs. M | UNFERTILIZED | 2015 | -1.92 | 0.518 | 10 | -3.71 | 0.010 | 9.88 | -19.5 |
| M vs U | FERTILIZED | 2012 | -1.9 | 0.97 | 10 | -1.96 | 0.173 | 13.2 | - |
| P vs U | FERTILIZED | 2012 | -3.78 | 0.956 | 10 | -3.95 | 0.007 | 13.2 | -28.5 |
| P vs. M | FERTILIZED | 2012 | -1.88 | 0.518 | 10 | -3.62 | 0.012 | 11.3 | -16.6 |
| M vs U | FERTILIZED | 2013 | -1.91 | 0.97 | 10 | -1.97 | 0.171 | 11 | - |
| P vs U | FERTILIZED | 2013 | -3.94 | 0.956 | 10 | -4.12 | 0.005 | 11 | -35.8 |
| P vs. M | FERTILIZED | 2013 | -2.03 | 0.518 | 10 | -3.91 | 0.007 | 9.08 | -22.3 |
| M vs U | FERTILIZED | 2014 | -0.318 | 0.97 | 10 | -0.327 | 0.943 | 9.11 | - |
| P vs U | FERTILIZED | 2014 | -1.99 | 0.956 | 10 | -2.08 | 0.144 | 9.11 | - |
| P vs. M | FERTILIZED | 2014 | -1.67 | 0.518 | 10 | -3.22 | 0.023 | 8.79 | -19 |
| M vs U | FERTILIZED | 2015 | 0.616 | 0.97 | 10 | 0.635 | 0.805 | 7.09 | - |
| P vs U | FERTILIZED | 2015 | -1.1 | 0.956 | 10 | -1.15 | 0.510 | 7.09 | - |
| P vs. M | FERTILIZED | 2015 | -1.71 | 0.518 | 10 | -3.3 | 0.020 | 7.7 | -22.2 |
| **Consumer pressure** | | | | | | | | | |
| Spraying | UNFERTILIZED | 2012 | 0.62 | 0.424 | 236 | 1.46 | 0.145 | 11.4 | - |
| Spraying | UNFERTILIZED | 2013 | -0.583 | 0.424 | 236 | -1.38 | 0.170 | 10.9 | - |
| Spraying | UNFERTILIZED | 2014 | -2.03 | 0.424 | 236 | -4.79 | 0.000 | 11.7 | -17.4 |
| Spraying | UNFERTILIZED | 2015 | -1.88 | 0.424 | 236 | -4.44 | 0.000 | 10.3 | -18.4 |
| Spraying | FERTILIZED | 2012 | -0.369 | 0.424 | 236 | -0.87 | 0.385 | 12 | - |
| Spraying | FERTILIZED | 2013 | -0.703 | 0.424 | 236 | -1.66 | 0.099 | 10.3 | - |
| Spraying | FERTILIZED | 2014 | -0.364 | 0.424 | 236 | -0.857 | 0.392 | 9.67 | - |
| Spraying | FERTILIZED | 2015 | -0.169 | 0.424 | 236 | -0.398 | 0.691 | 8.37 | - |
| **Soil resource supply** | | | | | | | | | |
| Fertilization | UNSPRAYED | 2012 | 0.104 | 0.425 | 236 | 0.244 | 0.807 | 11.4 | - |
| Fertilization | UNSPRAYED | 2013 | -1.52 | 0.425 | 236 | -3.57 | 0.000 | 10.9 | -13.9 |
| Fertilization | UNSPRAYED | 2014 | -3.18 | 0.425 | 236 | -7.48 | 0.000 | 11.7 | -27.2 |
| Fertilization | UNSPRAYED | 2015 | -3.25 | 0.425 | 236 | -7.64 | 0.000 | 10.3 | -31.7 |
| Fertilization | SPRAYED | 2012 | -0.886 | 0.425 | 236 | -2.09 | 0.038 | 12 | -7.36 |
| Fertilization | SPRAYED | 2013 | -1.64 | 0.425 | 236 | -3.85 | 0.000 | 10.3 | -15.8 |
| Fertilization | SPRAYED | 2014 | -1.51 | 0.425 | 236 | -3.55 | 0.000 | 9.67 | -15.6 |
| Fertilization | SPRAYED | 2015 | -1.53 | 0.425 | 236 | -3.61 | 0.000 | 8.37 | -18.3 |

**Table S5** Total species evenness model. Summary of most parsimonious interaction model of total species evenness in response to priority effects, consumer pressure, and soil resource supply across time.

| *Effect* | *DF (Num,Den)* | *F value* | *P value* |
|---:|---:|---:|---:|
| *(Intercept)* | 1,765 | 3349.9 | < 0.001 |
| *Priority* | 2,10 | 7.29 | 0.011 |
| *Consumers* | 1,236 | 0.47 | 0.492 |
| *Soil Resources* | 1,236 | 16.26 | < 0.001 |
| *Year* | 3,765 | 28.97 | < 0.001 |
| *Block* | 4,236 | 4.79 | 0.001 |
| *Priority:Consumers* | 2,236 | 0.61 | 0.546 |
| *Priority:Soil Resources* | 2,236 | 6.22 | 0.002 |
| *Consumers:Soil Resources* | 1,236 | 1.17 | 0.28 |
| *Priority:Year* | 6,765 | 8.04 | < 0.001 |
| *Consumers:Year* | 3,765 | 2.14 | 0.094 |
| *Soil Resources:Year* | 3,765 | 13.22 | < 0.001 |

**Table S6** Total species evenness multiple comparisons. Results of Least-Square Means post-hoc Tukey tests and multiple comparisons on total species evenness. The comparison shows the treatment vs. control, in the case of the priority effect treatment this compares the polyculture (P), monoculture (M), and unplanted (U). Interaction indicates whether the effect was in unfertilized plots or fertilized for priority effect and spraying treatments, and unsprayed or sprayed for the fertilization treatment. The estimated control is the model estimated value. For priority effect this shows the estimated mean value of the lower planted richness level. Estimated effect size is the estimated effect divided by estimated control for comparisons with a significant value ($p < 0.05$).

| Comparison | Interaction | Year | Estimated Effect | Standard Error | DF | T-ratio | p-value | Estimated Control* | Estimated effect size |
|---|---|---|---|---|---|---|---|---|---|
| | | | | **Priority Effects** | | | | | |
| M vs U | UNFERTILIZED | 2012 | -0.242 | 0.045 | 10 | -5.39 | 0.001 | 0.599 | -40.4 |
| P vs U | UNFERTILIZED | 2012 | -0.109 | 0.0442 | 10 | -2.45 | 0.080 | 0.599 | - |
| P vs. M | UNFERTILIZED | 2012 | 0.134 | 0.0238 | 10 | 5.61 | 0.001 | 0.357 | 37.5 |
| M vs U | UNFERTILIZED | 2013 | -0.0231 | 0.045 | 10 | -0.514 | 0.866 | 0.478 | - |
| P vs U | UNFERTILIZED | 2013 | 0.05 | 0.0442 | 10 | 1.13 | 0.518 | 0.478 | - |
| P vs. M | UNFERTILIZED | 2013 | 0.0732 | 0.0238 | 10 | 3.07 | 0.029 | 0.454 | 16.1 |
| M vs U | UNFERTILIZED | 2014 | -0.0626 | 0.045 | 10 | -1.39 | 0.381 | 0.548 | - |
| P vs U | UNFERTILIZED | 2014 | 0.00559 | 0.0442 | 10 | 0.126 | 0.991 | 0.548 | - |
| P vs. M | UNFERTILIZED | 2014 | 0.0682 | 0.0238 | 10 | 2.86 | 0.041 | 0.486 | 14 |
| M vs U | UNFERTILIZED | 2015 | -0.0175 | 0.045 | 10 | -0.389 | 0.920 | 0.511 | - |
| P vs U | UNFERTILIZED | 2015 | 0.0327 | 0.0442 | 10 | 0.74 | 0.746 | 0.511 | - |
| P vs. M | UNFERTILIZED | 2015 | 0.0502 | 0.0238 | 10 | 2.11 | 0.138 | 0.494 | - |
| M vs U | FERTILIZED | 2012 | -0.131 | 0.045 | 10 | -2.92 | 0.038 | 0.547 | -24 |
| P vs U | FERTILIZED | 2012 | -0.0215 | 0.0442 | 10 | -0.487 | 0.879 | 0.547 | - |
| P vs. M | FERTILIZED | 2012 | 0.11 | 0.0238 | 10 | 4.6 | 0.003 | 0.416 | 26.3 |
| M vs U | FERTILIZED | 2013 | 0.0881 | 0.045 | 10 | 1.96 | 0.173 | 0.478 | - |
| P vs U | FERTILIZED | 2013 | 0.137 | 0.0442 | 10 | 3.1 | 0.028 | 0.478 | 28.7 |
| P vs. M | FERTILIZED | 2013 | 0.049 | 0.0238 | 10 | 2.06 | 0.149 | 0.566 | - |
| M vs U | FERTILIZED | 2014 | 0.0486 | 0.045 | 10 | 1.08 | 0.547 | 0.501 | - |
| P vs U | FERTILIZED | 2014 | 0.0926 | 0.0442 | 10 | 2.09 | 0.141 | 0.501 | - |
| P vs. M | FERTILIZED | 2014 | 0.0441 | 0.0238 | 10 | 1.85 | 0.204 | 0.55 | - |
| M vs U | FERTILIZED | 2015 | 0.0937 | 0.045 | 10 | 2.08 | 0.143 | 0.387 | - |
| P vs U | FERTILIZED | 2015 | 0.12 | 0.0442 | 10 | 2.71 | 0.053 | 0.387 | - |
| P vs. M | FERTILIZED | 2015 | 0.0261 | 0.0238 | 10 | 1.09 | 0.539 | 0.48 | - |
| | | | | | **Consumer pressure** | | | | |
| Spraying | UNFERTILIZED | 2012 | -0.0175 | 0.0181 | 236 | -0.965 | 0.336 | 0.491 | - |
| Spraying | UNFERTILIZED | 2013 | -0.00995 | 0.0181 | 236 | -0.549 | 0.584 | 0.492 | - |
| Spraying | UNFERTILIZED | 2014 | 0.0214 | 0.0181 | 236 | 1.180 | 0.239 | 0.518 | - |
| Spraying | UNFERTILIZED | 2015 | 0.0249 | 0.0181 | 236 | 1.370 | 0.171 | 0.504 | - |
| Spraying | FERTILIZED | 2012 | -0.00051 | 0.0181 | 236 | -0.028 | 0.977 | 0.473 | - |
| Spraying | FERTILIZED | 2013 | 0.00703 | 0.0181 | 236 | 0.388 | 0.699 | 0.482 | - |
| Spraying | FERTILIZED | 2014 | 0.0384 | 0.0181 | 236 | 2.120 | 0.035 | 0.54 | 7.11 |
| Spraying | FERTILIZED | 2015 | 0.0418 | 0.0181 | 236 | 2.310 | 0.022 | 0.529 | 7.92 |
| | | | | | **Soil resource supply** | | | | |
| Fertilization | UNSPRAYED | 2012 | 0.006 | 0.0181 | 236 | 0.306 | 0.760 | 0.491 | - |
| Fertilization | UNSPRAYED | 2013 | 0.058 | 0.0181 | 236 | 3.220 | 0.001 | 0.492 | 11.9 |
| Fertilization | UNSPRAYED | 2014 | 0.010 | 0.0181 | 236 | 0.575 | 0.566 | 0.518 | - |
| Fertilization | UNSPRAYED | 2015 | -0.067 | 0.0181 | 236 | -3.690 | 0.000 | 0.504 | -13.3 |
| Fertilization | SPRAYED | 2012 | 0.023 | 0.0181 | 236 | 1.240 | 0.215 | 0.473 | - |
| Fertilization | SPRAYED | 2013 | 0.075 | 0.0181 | 236 | 4.160 | 0.000 | 0.482 | 15.7 |
| Fertilization | SPRAYED | 2014 | 0.027 | 0.0181 | 236 | 1.510 | 0.132 | 0.540 | - |
| Fertilization | SPRAYED | 2015 | -0.050 | 0.0181 | 236 | -2.750 | 0.006 | 0.529 | -9.43 |

**Table S7** Colonizing species evenness model. Summary of most parsimonious interaction model of colonizing species evenness in response to priority effects, consumer pressure, and soil resource supply across time.

| *Effect* | *DF (Num,Den)* | *F value* | *P value* |
|---|---|---|---|
| *(Intercept)* | 1,765 | 12689.66 | < 0.001 |
| *Priority* | 2,10 | 5.16 | 0.029 |
| *Consumers* | 1,236 | 5.99 | 0.015 |
| *Soil Resources* | 1,236 | 6.42 | 0.012 |
| *Year* | 3,765 | 81.03 | < 0.001 |
| *Block* | 4,236 | 5.73 | < 0.001 |
| *Priority:Consumers* | 2,236 | 1.64 | 0.197 |
| *Priority:Soil Resources* | 2,236 | 0.92 | 0.398 |
| *Consumers:Soil Resources* | 1,236 | 0.35 | 0.555 |
| *Priority:Year* | 6,765 | 0.4 | 0.881 |
| *Consumers:Year* | 3,765 | 2.84 | 0.037 |
| *Soil Resources:Year* | 3,765 | 0.93 | 0.426 |

**Table S8** Colonizing species evenness multiple comparisons. Results of Least-Square Means post-hoc Tukey tests and multiple comparisons on colonizing species evenness. The comparison shows the treatment vs. control, in the case of the priority effect treatment this compares the polyculture (P), monoculture (M), and unplanted (U). Interaction indicates whether the effect was in unfertilized plots or fertilized for priority effect and spraying treatments, and unsprayed or sprayed for the fertilization treatment. The estimated control is the model estimated value. For priority effect this shows the estimated mean value of the lower planted richness level. Estimated effect size is the estimated effect divided by estimated control for comparisons with a significant value ($p < 0.05$).

| Comparison | Interaction | Year | Estimated Effect | Standard Error | DF | T-ratio | p-value | Estimated Control* | Estimated effect size |
|---|---|---|---|---|---|---|---|---|---|
| **Priority Effects** | | | | | | | | | |
| M vs U | UNFERTILIZED | 2012 | 0.040 | 0.038 | 10 | 1.05 | 0.561 | 0.617 | - |
| P vs U | UNFERTILIZED | 2012 | 0.101 | 0.038 | 10 | 2.65 | 0.058 | 0.617 | - |
| P vs. M | UNFERTILIZED | 2012 | 0.060 | 0.022 | 10 | 2.81 | 0.045 | 0.658 | 9.19 |
| M vs U | UNFERTILIZED | 2013 | 0.078 | 0.038 | 10 | 2.03 | 0.155 | 0.496 | - |
| P vs U | UNFERTILIZED | 2013 | 0.163 | 0.038 | 10 | 4.29 | 0.004 | 0.496 | 32.8 |
| P vs. M | UNFERTILIZED | 2013 | 0.085 | 0.022 | 10 | 3.96 | 0.007 | 0.574 | 14.9 |
| M vs U | UNFERTILIZED | 2014 | -0.018 | 0.038 | 10 | -0.467 | 0.888 | 0.542 | - |
| P vs U | UNFERTILIZED | 2014 | 0.035 | 0.038 | 10 | 0.912 | 0.646 | 0.542 | - |
| P vs. M | UNFERTILIZED | 2014 | 0.052 | 0.022 | 10 | 2.43 | 0.082 | 0.525 | - |
| M vs U | UNFERTILIZED | 2015 | 0.010 | 0.038 | 10 | 0.25 | 0.966 | 0.482 | - |
| P vs U | UNFERTILIZED | 2015 | 0.045 | 0.038 | 10 | 1.18 | 0.490 | 0.482 | - |
| P vs. M | UNFERTILIZED | 2015 | 0.035 | 0.022 | 10 | 1.64 | 0.275 | 0.492 | - |
| M vs U | FERTILIZED | 2012 | 0.109 | 0.038 | 10 | 2.86 | 0.041 | 0.529 | 20.6 |
| P vs U | FERTILIZED | 2012 | 0.142 | 0.038 | 10 | 3.75 | 0.010 | 0.529 | 26.9 |
| P vs. M | FERTILIZED | 2012 | 0.033 | 0.022 | 10 | 1.54 | 0.313 | 0.638 | - |
| M vs U | FERTILIZED | 2013 | 0.146 | 0.038 | 10 | 3.84 | 0.008 | 0.459 | 31.9 |
| P vs U | FERTILIZED | 2013 | 0.204 | 0.038 | 10 | 5.39 | 0.001 | 0.459 | 44.5 |
| P vs. M | FERTILIZED | 2013 | 0.058 | 0.022 | 10 | 2.7 | 0.054 | 0.606 | - |
| M vs U | FERTILIZED | 2014 | 0.051 | 0.038 | 10 | 1.34 | 0.407 | 0.507 | - |
| P vs U | FERTILIZED | 2014 | 0.076 | 0.038 | 10 | 2.01 | 0.160 | 0.507 | - |
| P vs. M | FERTILIZED | 2014 | 0.025 | 0.022 | 10 | 1.17 | 0.496 | 0.558 | - |
| M vs U | FERTILIZED | 2015 | 0.079 | 0.038 | 10 | 2.06 | 0.149 | 0.416 | - |
| P vs U | FERTILIZED | 2015 | 0.087 | 0.038 | 10 | 2.28 | 0.105 | 0.416 | - |
| P vs. M | FERTILIZED | 2015 | 0.008 | 0.022 | 10 | 0.374 | 0.926 | 0.494 | - |
| **Consumer pressure** | | | | | | | | | |
| Spraying | UNFERTILIZED | 2012 | -0.011 | 0.020 | 236 | -0.533 | 0.595 | 0.67 | - |
| Spraying | UNFERTILIZED | 2013 | 0.027 | 0.020 | 236 | 1.35 | 0.178 | 0.563 | - |
| Spraying | UNFERTILIZED | 2014 | 0.044 | 0.020 | 236 | 2.25 | 0.025 | 0.526 | 8.43 |
| Spraying | UNFERTILIZED | 2015 | 0.037 | 0.020 | 236 | 1.89 | 0.061 | 0.482 | - |
| Spraying | FERTILIZED | 2012 | -0.003 | 0.020 | 236 | -0.169 | 0.866 | 0.659 | - |
| Spraying | FERTILIZED | 2013 | 0.034 | 0.020 | 236 | 1.71 | 0.088 | 0.59 | - |
| Spraying | FERTILIZED | 2014 | 0.052 | 0.020 | 236 | 2.61 | 0.010 | 0.57 | 9.03 |
| Spraying | FERTILIZED | 2015 | 0.044 | 0.020 | 236 | 2.25 | 0.025 | 0.519 | 8.54 |
| **Soil resource supply** | | | | | | | | | |
| Fertilization | UNSPRAYED | 2012 | -0.055 | 0.020 | 236 | -2.79 | 0.006 | 0.67 | -8.22 |
| Fertilization | UNSPRAYED | 2013 | -0.004 | 0.020 | 236 | -0.184 | 0.854 | 0.563 | - |
| Fertilization | UNSPRAYED | 2014 | -0.002 | 0.020 | 236 | -0.11 | 0.913 | 0.526 | - |
| Fertilization | UNSPRAYED | 2015 | -0.034 | 0.020 | 236 | -1.7 | 0.091 | 0.482 | - |
| Fertilization | SPRAYED | 2012 | -0.048 | 0.020 | 236 | -2.43 | 0.016 | 0.659 | -7.26 |
| Fertilization | SPRAYED | 2013 | 0.004 | 0.020 | 236 | 0.18 | 0.857 | 0.59 | - |
| Fertilization | SPRAYED | 2014 | 0.005 | 0.020 | 236 | 0.254 | 0.800 | 0.57 | - |
| Fertilization | SPRAYED | 2015 | -0.026 | 0.020 | 236 | -1.34 | 0.183 | 0.519 | - |

**Table S9** CWM of colonizing species' seed mass model. Summary of most parsimonious interaction model of CWM of colonizing species' seed mass in response to priority effects, consumer pressure, and soil resource supply across time.

| *Effect* | *DF (Num,Den)* | *F value* | *P value* |
|---|---|---|---|
| *(Intercept)* | 1,765 | 7.34 | 0.007 |
| *Priority* | 2,10 | 4.75 | 0.036 |
| *Consumers* | 1,236 | 0.04 | 0.846 |
| *Soil Resources* | 1,236 | 17.19 | < 0.001 |
| *Year* | 3,765 | 258.64 | < 0.001 |
| *Block* | 4,236 | 3.83 | 0.005 |
| *Priority:Consumers* | 2,236 | 1.2 | 0.304 |
| *Priority:Soil Resources* | 2,236 | 6.39 | 0.002 |
| *Consumers:Soil Resources* | 1,236 | 2.08 | 0.15 |
| *Priority:Year* | 6,765 | 2.23 | 0.039 |
| *Consumers:Year* | 3,765 | 1.8 | 0.146 |
| *Soil Resources:Year* | 3,765 | 9.92 | < 0.001 |

**Table S10** CWM of colonizing species' seed mass multiple comparisons. Results of Least-Square Means post-hoc Tukey tests and multiple comparisons on CWM of colonizing species' seed mass. The comparison shows the treatment vs. control, in the case of the priority effect treatment this compares the polyculture (P), monoculture (M), and unplanted (U). Interaction indicates whether the effect was in unfertilized plots or fertilized for priority effect and spraying treatments, and unsprayed or sprayed for the fertilization treatment. The estimated control is the model estimated value. For priority effect this shows the estimated mean value of the lower planted richness level. Estimated effect size is the back-transformed estimated effect plus estimated control divided by the back-transformed estimated control for comparisons with a significant value ($p < 0.05$).

| Comparison | Interaction | Year | Estimated Effect | Standard Error | DF | T-ratio | p-value | Estimated Control* | Estimated effect size |
|---|---|---|---|---|---|---|---|---|---|
| **Priority Effects** | | | | | | | | | |
| M vs U | UNFERTILIZED | 2012 | 0.677 | 0.184 | 10 | 3.67 | 0.011 | -1.16 | 96.8 |
| P vs U | UNFERTILIZED | 2012 | 0.891 | 0.186 | 10 | 4.78 | 0.002 | -1.16 | 144 |
| P vs. M | UNFERTILIZED | 2012 | 0.214 | 0.106 | 10 | 2.01 | 0.159 | -0.478 | - |
| M vs U | UNFERTILIZED | 2013 | 0.398 | 0.184 | 10 | 2.16 | 0.128 | -1.03 | - |
| P vs U | UNFERTILIZED | 2013 | 0.676 | 0.186 | 10 | 3.63 | 0.012 | -1.03 | 96.7 |
| P vs. M | UNFERTILIZED | 2013 | 0.279 | 0.106 | 10 | 2.62 | 0.061 | -0.631 | - |
| M vs U | UNFERTILIZED | 2014 | 0.283 | 0.184 | 10 | 1.53 | 0.318 | -0.613 | - |
| P vs U | UNFERTILIZED | 2014 | 0.545 | 0.186 | 10 | 2.92 | 0.037 | -0.613 | 72.4 |
| P vs. M | UNFERTILIZED | 2014 | 0.262 | 0.106 | 10 | 2.46 | 0.078 | -0.33 | - |
| M vs U | UNFERTILIZED | 2015 | 0.239 | 0.184 | 10 | 1.29 | 0.430 | 0.0585 | - |
| P vs U | UNFERTILIZED | 2015 | 0.495 | 0.186 | 10 | 2.65 | 0.058 | 0.0585 | - |
| P vs. M | UNFERTILIZED | 2015 | 0.256 | 0.106 | 10 | 2.41 | 0.086 | 0.297 | - |
| M vs U | FERTILIZED | 2012 | 0.342 | 0.184 | 10 | 1.85 | 0.203 | -0.866 | - |
| P vs U | FERTILIZED | 2012 | 0.423 | 0.186 | 10 | 2.27 | 0.107 | -0.866 | - |
| P vs. M | FERTILIZED | 2012 | 0.081 | 0.106 | 10 | 0.76 | 0.735 | -0.524 | - |
| M vs U | FERTILIZED | 2013 | 0.063 | 0.184 | 10 | 0.339 | 0.939 | -0.534 | - |
| P vs U | FERTILIZED | 2013 | 0.208 | 0.186 | 10 | 1.11 | 0.527 | -0.534 | - |
| P vs. M | FERTILIZED | 2013 | 0.145 | 0.106 | 10 | 1.37 | 0.393 | -0.471 | - |
| M vs U | FERTILIZED | 2014 | -0.053 | 0.184 | 10 | -0.285 | 0.956 | 0.101 | - |
| P vs U | FERTILIZED | 2014 | 0.076 | 0.186 | 10 | 0.408 | 0.913 | 0.101 | - |
| P vs. M | FERTILIZED | 2014 | 0.129 | 0.106 | 10 | 1.21 | 0.474 | 0.0488 | - |
| M vs U | FERTILIZED | 2015 | -0.097 | 0.184 | 10 | -0.525 | 0.861 | 0.644 | - |
| P vs U | FERTILIZED | 2015 | 0.026 | 0.186 | 10 | 0.139 | 0.989 | 0.644 | - |
| P vs. M | FERTILIZED | 2015 | 0.123 | 0.106 | 10 | 1.15 | 0.505 | 0.548 | - |
| **Consumer pressure** | | | | | | | | | |
| Spraying | UNFERTILIZED | 2012 | 0.097 | 0.076 | 236 | 1.27 | 0.204 | -0.681 | - |
| Spraying | UNFERTILIZED | 2013 | -0.037 | 0.076 | 236 | -0.487 | 0.626 | -0.652 | - |
| Spraying | UNFERTILIZED | 2014 | -0.066 | 0.076 | 236 | -0.869 | 0.386 | -0.304 | - |
| Spraying | UNFERTILIZED | 2015 | 0.018 | 0.076 | 236 | 0.233 | 0.816 | 0.294 | - |
| Spraying | FERTILIZED | 2012 | -0.012 | 0.076 | 236 | -0.154 | 0.878 | -0.584 | - |
| Spraying | FERTILIZED | 2013 | -0.146 | 0.076 | 236 | -1.92 | 0.057 | -0.69 | - |
| Spraying | FERTILIZED | 2014 | -0.175 | 0.076 | 236 | -2.3 | 0.023 | -0.37 | -16.1 |
| Spraying | FERTILIZED | 2015 | -0.091 | 0.076 | 236 | -1.19 | 0.233 | 0.312 | - |
| **Soil resource supply** | | | | | | | | | |
| Fertilization | UNSPRAYED | 2012 | 0.076 | 0.076 | 236 | 0.998 | 0.319 | -0.681 | - |
| Fertilization | UNSPRAYED | 2013 | 0.282 | 0.076 | 236 | 3.7 | 0.000 | -0.652 | 32.6 |
| Fertilization | UNSPRAYED | 2014 | 0.501 | 0.076 | 236 | 6.57 | 0.000 | -0.304 | 65 |
| Fertilization | UNSPRAYED | 2015 | 0.372 | 0.076 | 236 | 4.89 | 0.000 | 0.294 | 45.1 |
| Fertilization | SPRAYED | 2012 | -0.033 | 0.076 | 236 | -0.43 | 0.668 | -0.584 | - |
| Fertilization | SPRAYED | 2013 | 0.173 | 0.076 | 236 | 2.27 | 0.024 | -0.69 | 18.9 |
| Fertilization | SPRAYED | 2014 | 0.392 | 0.076 | 236 | 5.14 | 0.000 | -0.37 | 48 |
| Fertilization | SPRAYED | 2015 | 0.263 | 0.076 | 236 | 3.46 | 0.001 | 0.312 | 30.1 |

**Table S11** Interspecific height model. Summary of most parsimonious interaction model interspecific height in response to priority effects, consumer pressure, and soil resource supply in the third year of study (2014).

| *Effect* | *DF (Num,Den)* | *F value* | *P value* |
|---|---|---|---|
| *(Intercept)* | 1,244 | 1802.92 | < 0.001 |
| *Priority* | 2,10 | 1.24 | 0.33 |
| *Consumers* | 1,244 | 2.11 | 0.147 |
| *Soil Resources* | 1,244 | 53.68 | < 0.001 |
| *Block* | 1,244 | 2.39 | 0.123 |

**Table S12** Intraspecific height model. Summary of most parsimonious interaction model intraspecific height in response to priority effects, consumer pressure, and soil resource supply in the third year of study (2014).

| *Effect* | *DF (Num,Den)* | *F value* | *P value* |
|---|---|---|---|
| *(Intercept)* | 1,239 | 29.31 | < 0.001 |
| *Priority* | 2,10 | 1.38 | 0.296 |
| *Consumers* | 1,239 | 26.99 | < 0.001 |
| *Soil Resources* | 1,239 | 27.24 | < 0.001 |
| *Block* | 1,239 | 13.16 | < 0.001 |
| *Consumers:Soil Resources* | 1,239 | 4.28 | 0.04 |

**Table S13** Interspecific SLA model. Summary of most parsimonious interaction model interspecific SLA in response to priority effects, consumer pressure, and soil resource supply in the third year of study (2014).

| *Effect* | *DF (Num,Den)* | *F value* | *P value* |
|---|---|---|---|
| *(Intercept)* | 1,244 | 4809.64 | < 0.001 |
| *Priority* | 2,10 | 0.81 | 0.474 |
| *Consumers* | 1,244 | 11.79 | 0.001 |
| *Soil Resources* | 1,244 | 11.94 | 0.001 |
| *Block* | 1,244 | 16.04 | < 0.001 |

**Table S14** Intraspecific SLA model. Summary of most parsimonious interaction model intraspecific SLA in response to priority effects, consumer pressure, and soil resource supply in the third year of study (2014).

| *Effect* | *DF (Num,Den)* | *F value* | *P value* |
|---|---|---|---|
| *(Intercept)* | 1,244 | 2 | 0.159 |
| *Priority* | 2,10 | 1.38 | 0.297 |
| *Consumers* | 1,244 | 0.64 | 0.423 |
| *Soil Resources* | 1,244 | 2.91 | 0.089 |
| *Block* | 1,244 | 23.8 | < 0.001 |

**Table S15** Light attenuation model. Summary of most parsimonious interaction model of light attenuation in response to priority effects, consumer pressure, and soil resource supply across time.

| *Effect* | *DF (Num,Den)* | *F value* | *P value* |
|---:|---:|---:|---:|
| *(Intercept)* | 1,753 | 806.35 | < 0.001 |
| *Priority* | 2,10 | 1.71 | 0.23 |
| *Consumers* | 1,236 | 58.99 | < 0.001 |
| *Soil Resources* | 1,236 | 16.97 | < 0.001 |
| *Year* | 3,753 | 284.05 | < 0.001 |
| *Block* | 4,236 | 1.04 | 0.386 |
| *Priority:Consumers* | 2,236 | 0.33 | 0.719 |
| *Priority:Soil Resources* | 2,236 | 0.38 | 0.685 |
| *Consumers:Soil Resources* | 1,236 | 15.21 | < 0.001 |
| *Priority:Year* | 6,753 | 36 | < 0.001 |
| *Consumers:Year* | 3,753 | 14.74 | < 0.001 |
| *Soil Resources:Year* | 3,753 | 27.46 | < 0.001 |
| *Priority:Consumers:Year* | 6,753 | 2.42 | 0.025 |
| *Priority:Soil Resources:Year* | 6,753 | 2.25 | 0.037 |

**Table S16** Light attenuation multiple comparisons. Results of Least-Square Means post-hoc Tukey tests and multiple comparisons on light attenuation. The comparison shows the treatment vs. control, in the case of the priority effect treatment this compares the polyculture (P), monoculture (M), and unplanted (U). Interaction indicates whether the effect was in unfertilized plots or fertilized for priority effect and spraying treatments, and unsprayed or sprayed for the fertilization treatment. The estimated control is the model estimated value. For priority effect this shows the estimated mean value of the lower planted richness level. Estimated effect size is the estimated effect divided by estimated control for comparisons with a significant value ($p < 0.05$).

| Comparison | Interaction | Year | Estimated Effect | Standard Error | DF | T-ratio | p-value | Estimated Control* | Estimated effect size |
|---|---|---|---|---|---|---|---|---|---|
| | | | | **Priority Effects** | | | | | |
| M vs U | UNFERTILIZED | 2012 | -0.302 | 0.031 | 10 | -9.67 | 0.000 | 0.541 | -55.8 |
| P vs U | UNFERTILIZED | 2012 | -0.346 | 0.029 | 10 | -12.10 | 0.000 | 0.541 | -64 |
| P vs. M | UNFERTILIZED | 2012 | -0.044 | 0.021 | 10 | -2.08 | 0.143 | 0.239 | - |
| M vs U | UNFERTILIZED | 2013 | -0.003 | 0.008 | 10 | -0.40 | 0.917 | 0.048 | - |
| P vs U | UNFERTILIZED | 2013 | 0.002 | 0.007 | 10 | 0.23 | 0.972 | 0.048 | - |
| P vs. M | UNFERTILIZED | 2013 | 0.005 | 0.005 | 10 | 0.89 | 0.661 | 0.045 | - |
| M vs U | UNFERTILIZED | 2014 | 0.006 | 0.003 | 10 | 1.75 | 0.234 | 0.012 | - |
| P vs U | UNFERTILIZED | 2014 | 0.003 | 0.003 | 10 | 1.14 | 0.515 | 0.012 | - |
| P vs. M | UNFERTILIZED | 2014 | -0.002 | 0.002 | 10 | -1.05 | 0.567 | 0.018 | - |
| M vs U | UNFERTILIZED | 2015 | 0.008 | 0.004 | 10 | 1.83 | 0.208 | 0.012 | - |
| P vs U | UNFERTILIZED | 2015 | 0.008 | 0.004 | 10 | 2.05 | 0.152 | 0.012 | - |
| P vs. M | UNFERTILIZED | 2015 | 0.000 | 0.003 | 10 | 0.05 | 0.998 | 0.020 | - |
| M vs U | FERTILIZED | 2012 | -0.152 | 0.031 | 10 | -4.88 | 0.002 | 0.304 | -50.1 |
| P vs U | FERTILIZED | 2012 | -0.220 | 0.029 | 10 | -7.70 | 0.000 | 0.304 | -72.4 |
| P vs. M | FERTILIZED | 2012 | -0.068 | 0.021 | 10 | -3.16 | 0.025 | 0.151 | -44.6 |
| M vs U | FERTILIZED | 2013 | 0.000 | 0.008 | 10 | 0.04 | 0.999 | 0.027 | - |
| P vs U | FERTILIZED | 2013 | 0.008 | 0.007 | 10 | 1.13 | 0.521 | 0.027 | - |
| P vs. M | FERTILIZED | 2013 | 0.008 | 0.005 | 10 | 1.45 | 0.353 | 0.028 | - |
| M vs U | FERTILIZED | 2014 | 0.003 | 0.003 | 10 | 0.86 | 0.677 | 0.009 | - |
| P vs U | FERTILIZED | 2014 | 0.002 | 0.003 | 10 | 0.62 | 0.813 | 0.009 | - |
| P vs. M | FERTILIZED | 2014 | -0.001 | 0.002 | 10 | -0.43 | 0.905 | 0.012 | - |
| M vs U | FERTILIZED | 2015 | 0.001 | 0.004 | 10 | 0.29 | 0.956 | 0.014 | - |
| P vs U | FERTILIZED | 2015 | 0.002 | 0.004 | 10 | 0.58 | 0.835 | 0.014 | - |
| P vs. M | FERTILIZED | 2015 | 0.001 | 0.003 | 10 | 0.35 | 0.934 | 0.016 | - |
| | | | | | **Consumer pressure** | | | | |
| Spraying | UNFERTILIZED | 2012 | -0.115 | 0.016 | 236.000 | -7.23 | 0.000 | 0.383 | -30 |
| Spraying | UNFERTILIZED | 2013 | -0.026 | 0.004 | 236 | -6.23 | 0.000 | 0.0607 | -43.4 |
| Spraying | UNFERTILIZED | 2014 | -0.012 | 0.002 | 236 | -5.9 | 0.000 | 0.021 | -57.9 |
| Spraying | UNFERTILIZED | 2015 | -0.012 | 0.002 | 236 | -4.72 | 0.000 | 0.0229 | -51.1 |
| Spraying | FERTILIZED | 2012 | -0.105 | 0.016 | 236 | -6.65 | 0.000 | 0.268 | -39.3 |
| Spraying | FERTILIZED | 2013 | -0.017 | 0.004 | 236 | -4.05 | 0.000 | 0.0344 | -49.8 |
| Spraying | FERTILIZED | 2014 | -0.003 | 0.002 | 236 | -1.41 | 0.159 | 0.00884 | - |
| Spraying | FERTILIZED | 2015 | -0.002 | 0.002 | 236 | -0.987 | 0.325 | 0.0112 | - |
| | | | | | **Soil resource supply** | | | | |
| Fertilization | UNSPRAYED | 2012 | -0.150 | 0.016 | 236 | -9.48 | 0.000 | 0.383 | -39.3 |
| Fertilization | UNSPRAYED | 2013 | -0.022 | 0.004 | 236 | -5.17 | 0.000 | 0.061 | -36 |
| Fertilization | UNSPRAYED | 2014 | -0.009 | 0.002 | 236 | -4.32 | 0.000 | 0.021 | -42.4 |
| Fertilization | UNSPRAYED | 2015 | -0.006 | 0.002 | 236 | -2.44 | 0.015 | 0.023 | -26.5 |
| Fertilization | SPRAYED | 2012 | -0.141 | 0.016 | 236 | -8.9 | 0.000 | 0.268 | -52.6 |
| Fertilization | SPRAYED | 2013 | -0.013 | 0.004 | 236 | -2.99 | 0.003 | 0.034 | -36.8 |
| Fertilization | SPRAYED | 2014 | 0.000 | 0.002 | 236 | 0.167 | 0.868 | 0.009 | - |
| Fertilization | SPRAYED | 2015 | 0.003 | 0.002 | 236 | 1.28 | 0.200 | 0.011 | - |

**Fig. S1 Correlations between light attenuation and total species richness across time. In years where a significant correlation ($p < 0.05$) was detected, a blue line of best fit was added with the gray 95% confidence interval across the range of values measured.**

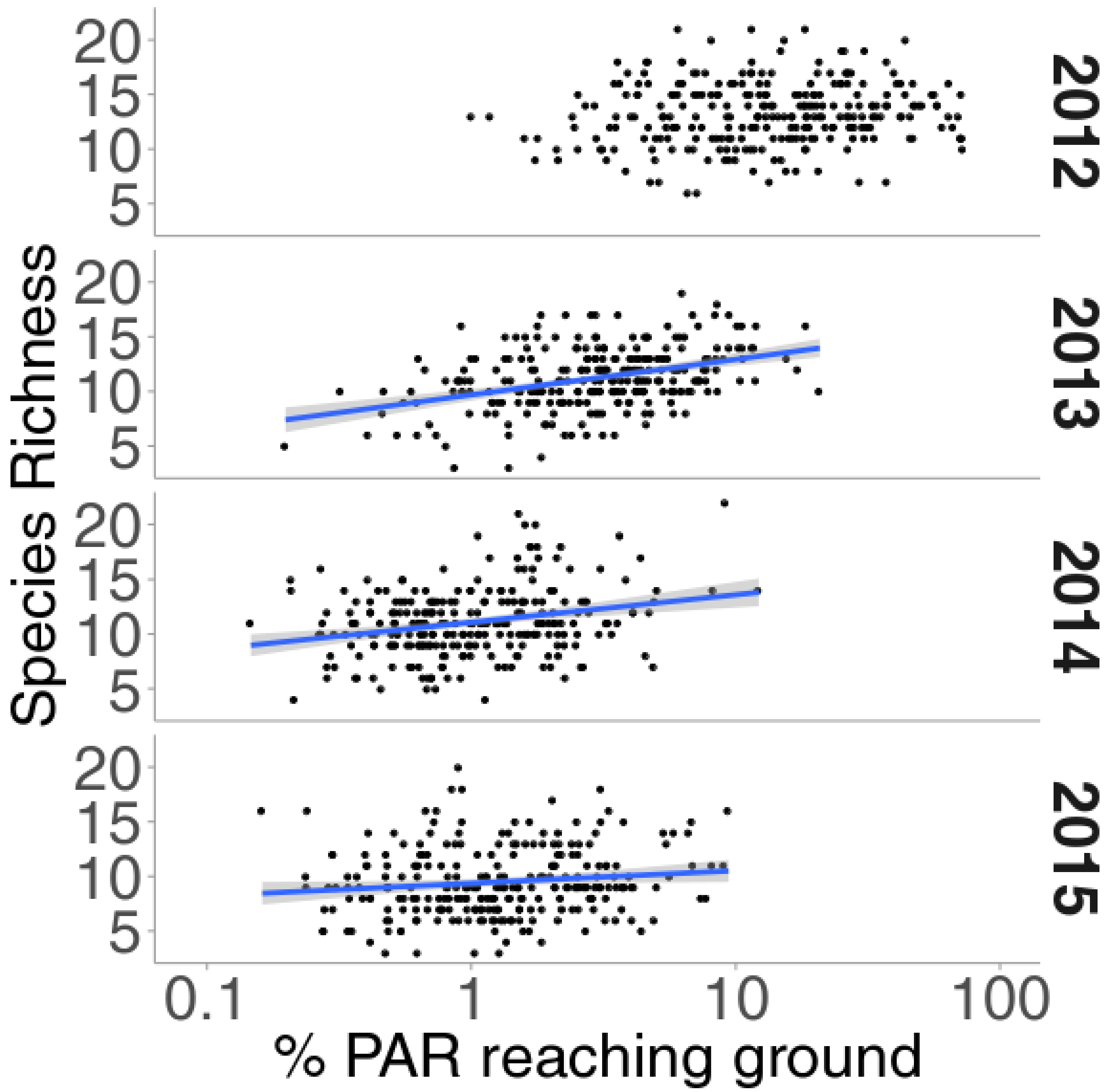

**Fig. S2 Correlations between light attenuation and total species evenness across time. In years where a significant correlation ($p < 0.05$) was detected, a blue line of best fit was added with the gray 95% confidence interval across the range of values measured.**

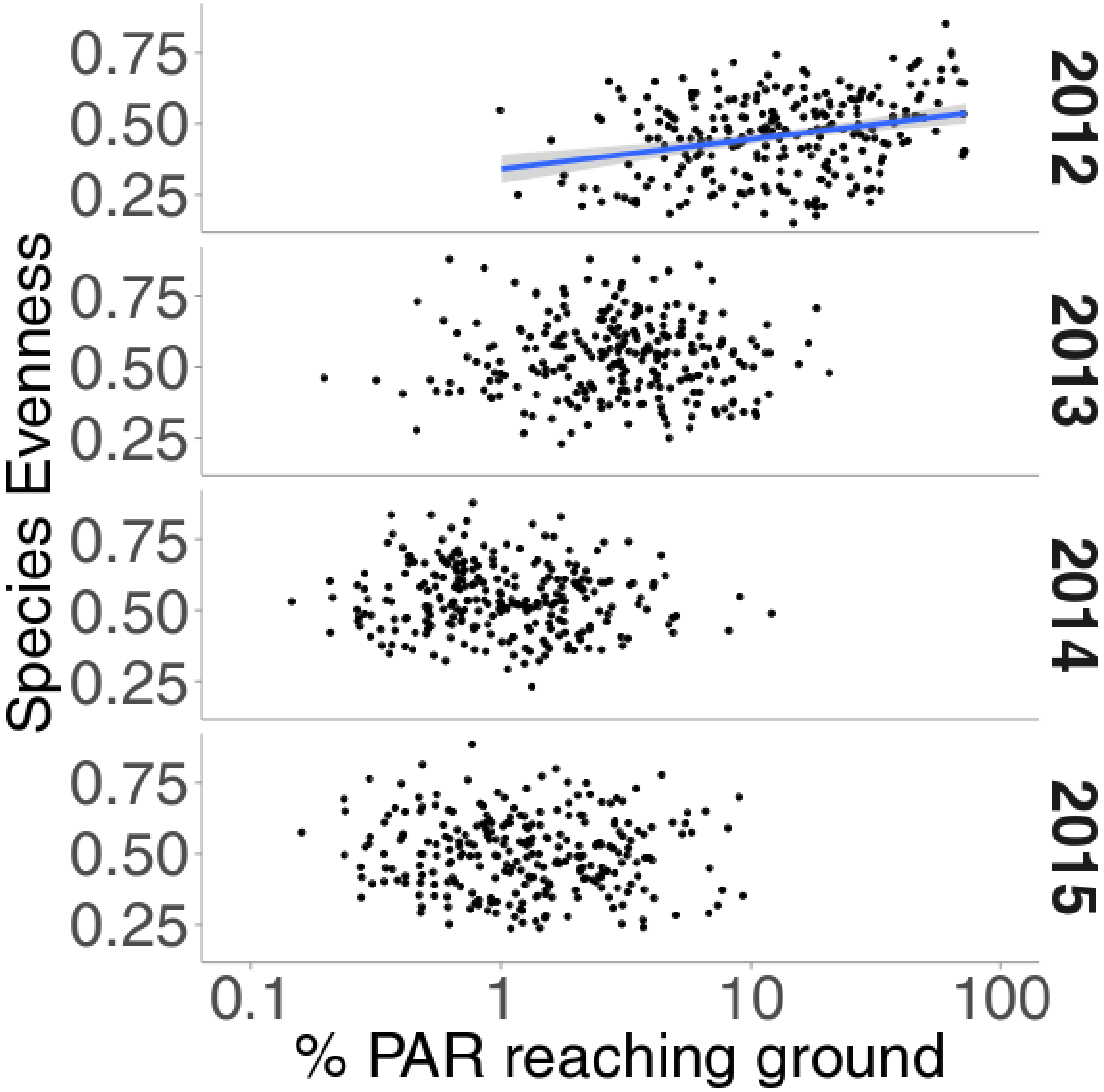

**Fig. S3 Correlations between light attenuation and colonizing species seed mass (CWM) across time. In years where a significant correlation ($p < 0.05$) was detected, a blue line of best fit was added with the gray 95% confidence interval across the range of values measured.**

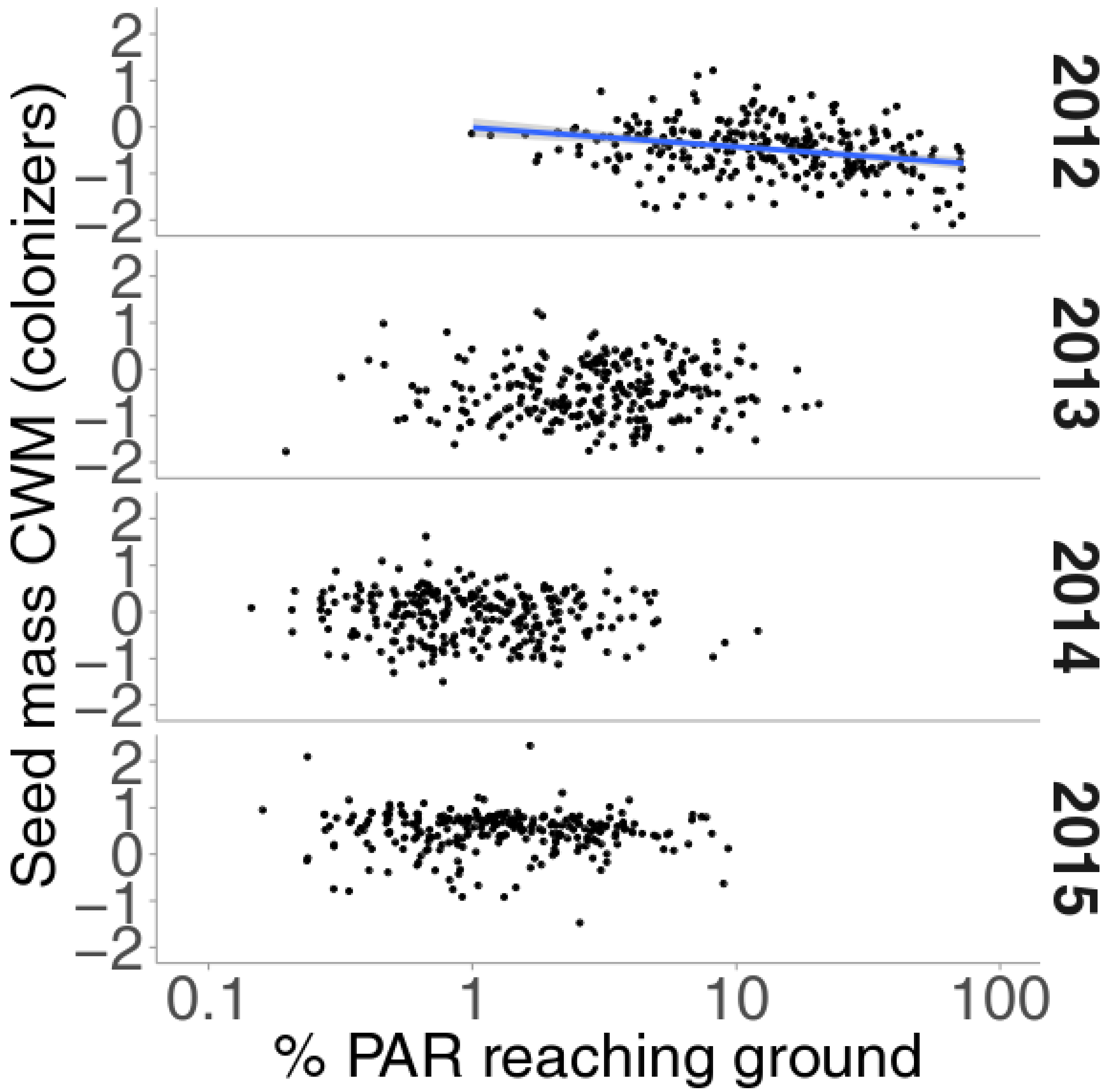

**Fig. S4 Correlation between light attenuation and interspecific height (CWM) in the third year of study (2014). A significant correlation ($p < 0.05$) was detected, a blue line of best fit was added with the gray 95% confidence interval across the range of values measured.**

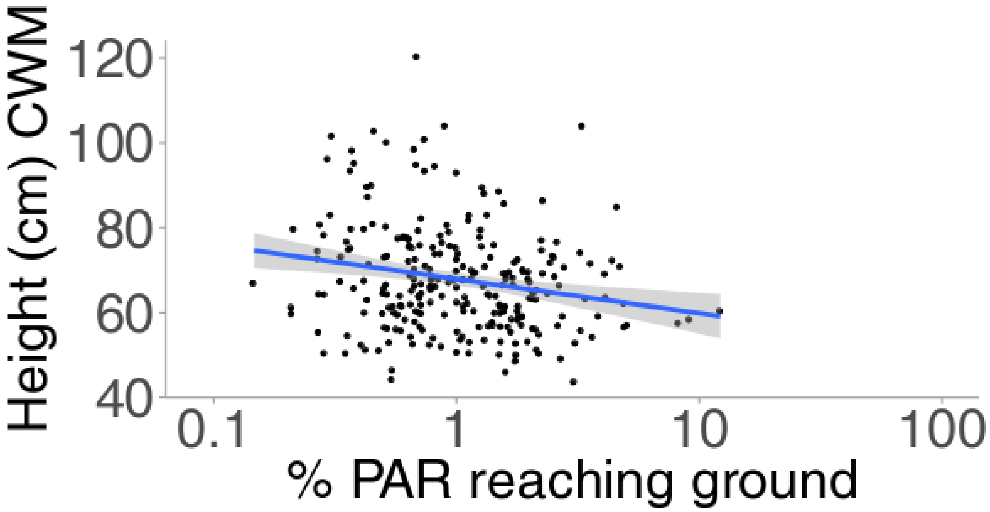


**Fig. S4 Correlation between light attenuation and interspecific SLA (CWM) in the third year of study (2014). No significant relationship was detected.**

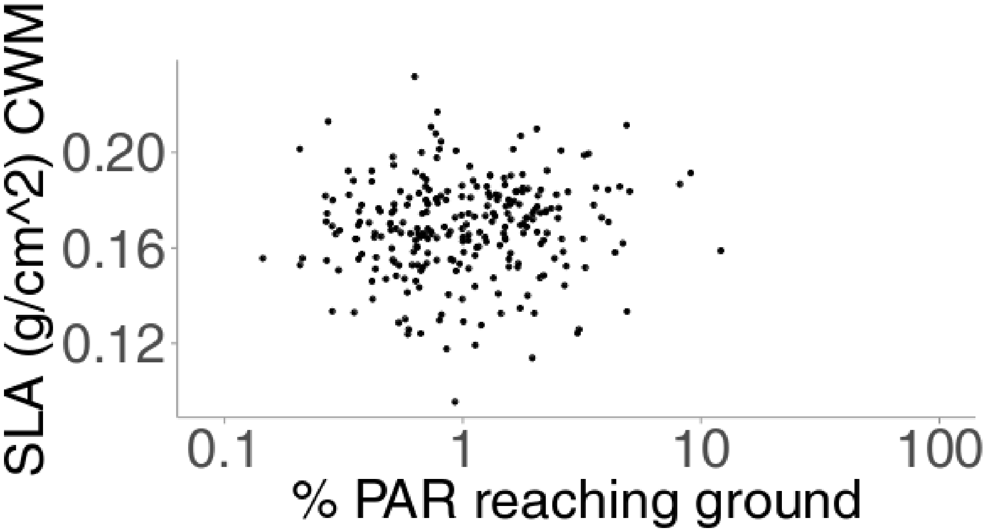
